# Molecular convergence by differential domain acquisition is a hallmark of chromosomal passenger complex evolution

**DOI:** 10.1101/2022.01.07.475321

**Authors:** Shinichiro Komaki, Eelco C. Tromer, Geert De Jaeger, Nancy De Winne, Maren Heese, Arp Schnittger

## Abstract

The chromosomal passenger complex (CPC) is a heterotetrameric regulator of eukaryotic cell division, consisting of an Aurora-type kinase and a scaffold built of INCENP, Borealin and Survivin. While most CPC components are conserved across eukaryotes, orthologs of the chromatin reader Survivin have previously only been found in animals and fungi, raising the question of how its essential role is carried out in other eukaryotes. By characterizing proteins that bind to the Arabidopsis Borealin ortholog, we identified BOREALIN RELATED INTERACTOR 1 and 2 (BORI1 and BORI2) as redundant Survivin-like proteins in the context of the CPC in plants. Loss of BORI function is lethal and a reduced expression of *BORIs* causes severe developmental defects. Similar to Survivin, we find that the BORIs bind to phosphorylated histone H3, relevant for correct CPC association with chromatin. However, this interaction is not mediated by a BIR domain as in previously recognized Survivin orthologs, but by an FHA domain, a widely conserved phosphate-binding module. We propose that the unifying criterion of Survivin-type proteins is a helix that facilitates complex formation with the other two scaffold components, and that the addition of a phosphate-binding domain, necessary for concentration at the inner centromere, evolved in parallel in different eukaryotic groups. Using sensitive similarity searches, we indeed find conservation of this helical domain between animals and plants, and identify the missing CPC component in most eukaryotic supergroups. Interestingly, we also detect Survivin orthologs without a defined phosphate-binding domain, possibly reflecting the situation in the last eukaryotic common ancestor.

**Significance Statement:** The identification of two *SURVIVIN*-type genes in the model plant Arabidopsis unfolded the evolutionary trajectories of this central chromosomal passenger complex component and led to the identification of SURVIVIN orthologs in almost the entire eukaryotic kingdom. Our work indicates that the central most aspect of the SURVIVIN gene family is a helix to make contact with two other core chromosomal passenger complex members whereas the addition of a phosphate-binding domain shown to bind to chromatin in animals and plants evolved in parallel at least 3 times in different eukaryotic branches.

## Introduction

Proper chromosome segregation and cytokinesis are essential for every organism to accurately transmit its genomic information to its progeny. For both processes, a precise regulation of the microtubule cytoskeleton is of key importance in plants and other eukaryotes (1). First, the microtubule fibers of the spindle have to be attached to chromosomes so that they will be equally distributed during cell division. The attachment is accomplished and monitored by a conserved large multi-protein structure called the kinetochore, which assembles at the centromeres of chromosomes (2, 3). Second, microtubules need to be precisely arranged to accomplish cytokinesis following chromosome segregation. In animals, microtubules mark the future site of division and facilitate the reorganization of actin and myosin at the cleavage furrow while in plants, microtubules form the scaffold of a cell wall generating structure at the plane of division, the phragmoplast (4).

A key regulator of microtubule organization is the multi-member Aurora kinase family that expanded through multiple independent parallel duplications in different eukaryotic clades (5). Aurora activity is linked to a protein assembly called chromosomal passenger complex (CPC) at both the centromeres and at the site of cytokinesis. The CPC is best studied in animals and yeast where besides at least one of the Aurora paralogs, it consists of three additional proteins: INCENP, Borealin, and Survivin. The C-terminus of INCENP contains a conserved motif, called IN-box, that directly binds to Aurora, while the N-terminus of INCENP forms a three-helical bundle with the C-termini of the other two CPC components Borealin and Survivin (6).

The localization of the CPC is highly dynamic during cell division and determines its function (6, 7). Before anaphase onset, the CPC localizes to the inner centromere where it monitors inter-kinetochore tension and prevents chromosome missegregation. During anaphase, the complex moves to the spindle midzone to promote cytokinesis (Fig. 1*A*). Multiple mechanisms have been shown to impinge on the correct localization of the CPC during mitosis. For instance, Borealin has been shown to confer a general affinity for nucleosomes required for chromosome association (8), but the CPC’s enrichment at centromeres in metaphase is dependent on at least two additional interconnected pathways (9). On the one hand, Borealin has been shown to interact with Shugoshin, which in turn is able to bind the centromere-associated histone mark H2AT120^ph^ deposited by the spindle checkpoint kinase Bub1. On the other hand Survivin binds via its Baculovirus IAP Repeat (BIR) domain to phosphorylated threonine 3 in the tail of histone H3 (H3T3^ph^) that becomes phosphorylated by the kinase Haspin (10–12). In addition, a recent study showed that Survivin is also able to bind Shugoshin at its N-terminus which resembles the phosphorylated H3-tail (13). Since both interactions involve the same site of Survivin’s BIR domain they are considered mutually exclusive which is in accordance with the model of two spatially distinct CPC pools: a Bub1-dependent kinetochore-proximal centromere pool, involving interactions of Survivin and Borealin with Shugoshin and a Haspin-dependent inner centromere pool entailing the H3T3^ph^ Survivin contact (14–16). The translocation of the CPC to the spindle midzone is then mediated by MKLP2, a member of the kinesin-6 family, which directly binds to the three-helical bundle of the CPC (17). However, this interaction only occurs during anaphase when a Cdk1-mediated inhibitory phosphorylation is removed from INCENP (18).

**Figure 1.**
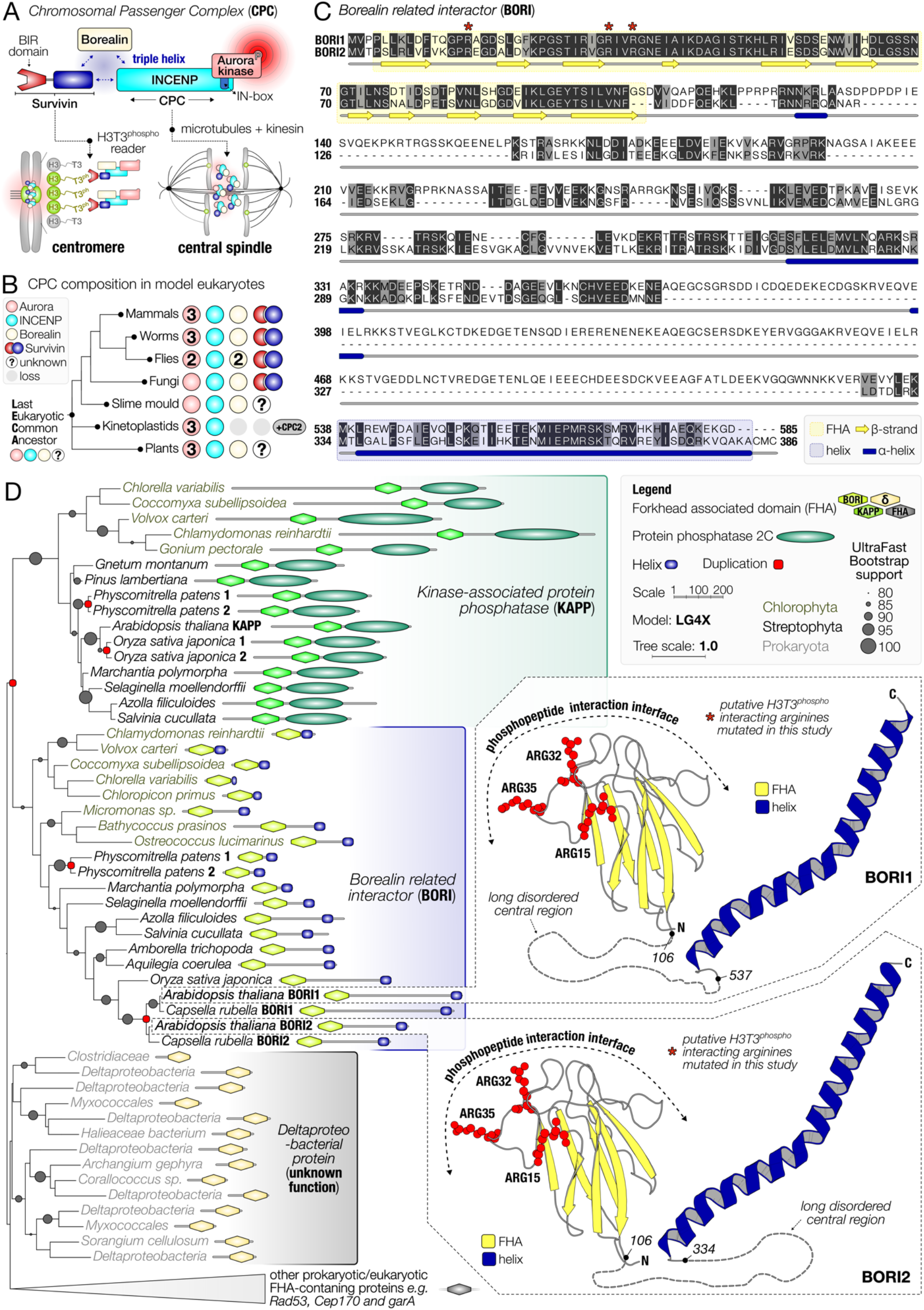
*BOREALIN RELATED INTERACTOR (BORI)* genes in plants. (**A**) The Chromosomal passenger complex (CPC) consists of an Aurora-type kinase scaffolded by the triple helix-based trimer INCENP, Borealin and Survivin. Metaphase CPC localization at the centromere is dependent on a Survivin-H3T3^ph^ interaction and anaphase localization at the central spindle relies on interactions with microtubules and kinesins (shown below). (**B**) Presence-absence matrix of CPC components in model organisms that have previously been found throughout the eukaryotic tree of life (2, 20). Colors of the proteins correspond to the cartoon in panel A. Numbers indicate the paralogs of a CPC component found in a particular clade. Question marks indicate the inability to detect orthologues. Greyed out circles indicate loss of components. (**C**) Multiple alignment of Borealin Related Interactor 1 and 2 found in *Arabidopsis thaliana*. FHA domains (yellow); C-terminal helices (dark blue). Secondary structure consensus of the Alphafold2 predicted 3D structures of BORI1 and 2 is projected below the alignment. Stars indicate three arginine residues (ARG-15-32-35), which likely face the phosphorylated histone H3 tail (see ball-and-sticks representation in panel D). Color scheme: 100% identity (black), similar physicochemical properties (grey), others (white). (**D**) Unrooted maximum-likelihood phylogenetic tree of FHA domains most similar to that of BORI orthologs, found amongst prokaryotes and eukaryotes. Domains are projected onto the phylogenetic tree and are to scale (see legend). Branch lengths are scaled and indicate the number of substitutions per site (see scale bar in the legend). Circles indicate bootstrap support (1000x replicates, only higher than 80% support shown). Red squares indicate duplication nodes. AthBORI1 and 2 Alphafold2-predicted 3D structures are shown on the right, with putative phosphate-interacting residues shown in a ball-and-sticks representation (see also panel C). Colors indicate the two different domains: FHA (yellow); helix (blue). Long disordered central regions are represented by dashed lines and are not to scale (see panel C for omitted residues). See *SI Appendix*, Dataset S2 for full-length 3D structures. For phylogenetic analysis details see *SI Appendix*, Dataset S3.

Comparative genomics and molecular analyses have revealed that Arabidopsis, and likely all other plants, are also equipped with a CPC similar to those found in animal and fungal model systems, which contains AUR3, one of three Aurora kinase paralogs in plants, INCENP, and a plant ortholog of Borealin, called BOREALIN RELATED (BORR) (2, 19–21). Loss of CPC function leads to gametophytic and sporophytic (embryonic) lethality in Arabidopsis underlining its key role in cell proliferation across the eukaryotic tree of life (19, 21). As seen in other eukaryotes, the plant CPC dynamically changes its subcellular localization throughout the cell cycle. It localizes to inner centromeres and prevents chromosome missegregation in early mitosis, and after anaphase onset it relocates to the center of the phragmoplast (21). Additionally, it was shown that plant Haspin phosphorylates histone H3 tails at threonine 3, and that this activity is needed to recruit AUR3 to the inner centromere (22–24). However, an ortholog of the H3T3^ph^-reader Survivin, needed for concentration of the CPC at the inner centromere, has not been identified in plants, nor any other eukaryotic lineages outside animals and fungi (20) (Fig. 1*B*).

Notably, in animals, next to its role in the CPC, Survivin also controls programmed cell death as a member of the inhibitor of apoptosis (IAP) protein family which is characterized by the presence of one to several BIR domains (25, 26). However, neither the IAP protein family, nor a bona fide BIR domain can be found in plants (27–29). Moreover, plants do not undergo apoptosis but display different mechanisms of programmed cell death instead (27–29).

In this study, we isolate BOREALIN-RELATED INTERACTOR 1 and 2 (BORI1 and 2) from Arabidopsis, and provide molecular and biochemical evidence that both proteins act as redundant plant analogs of Survivin with respect to its function in the CPC. Notably, instead of a BIR domain, BORIs contain an FHA domain to bind to phosphorylated histones. Furthermore, we reveal through extensive comparative genomics analyses that Survivin and BORI are indeed orthologs, and that the Survivin gene family is widely conserved amongst eukaryotes. The key characteristics of this Survivin/BORI gene family can be delineated to two functional domains with likely different evolutionary history: (I) a structurally conserved helix to make contact with the other subunits of the CPC, and (II) one domain characterized by convergent molecular evolution to mediate the interaction with H3T3^ph^ at the inner centromere.

## Results

### Identification of Borealin-interacting proteins in plants

Previously, we identified and functionally characterized a plant homolog of Borealin, called BOREALIN RELATED (BORR). BORR acts together with INCENP, also known as WYRD in *Arabidopsis thaliana (19)*, as the scaffold of the presumed equivalent of the Chromosomal Passenger Complex (CPC) in plants (21). However, an ortholog of Survivin, the third essential scaffolding subunit of the CPC, could so far not be detected outside animals and fungi (Fig. 1*B*), raising multiple different hypotheses on the absence of Survivin in plants and other eukaryotic lineages including: (I) lack of bioinformatic method sensitivity to detect Survivin orthologs, (II) existence of novel CPC subunits co-opting the function of Survivin, similar to CPC2 in the eukaryotic lineage Kinetoplastida (30), (III) coverage of its function by either BORR or WYRD (2, 20, 21), and (IV) lack of the necessity of Survivin function for the CPC in plants, i.e. Survivin representing an evolutionary novelty specific to animals and fungi (Fig. 1*B*).

To identify putative CPC-associated components in plants, we performed tandem affinity purification (TAP) followed by mass spectrometry (MS) using an Arabidopsis cell suspension culture expressing BORR with a CGSrhino tag at its C-terminus (31). The experiment was performed in duplicate and in both cases only one protein, At3g02400, passed all thresholds of the TAP evaluation pipeline and was subsequently named BOREALIN RELATED INTERACTOR 1 (BORI1). In addition, the known BORR interactor INCENP/WYRD was found in both experiments although each time only with one peptide, i.e., below the two-peptide cut-off of the standard evaluation pipeline (*SI Appendix*, Dataset S1).

At3g02400 was previously described as FORKHEAD-ASSOCIATED DOMAIN PROTEIN 3 (FHA3) and found to bind *in vitro* to a promoter fragment of PEROXIN 11b (PEX11b), which encodes a peroxisome protein. It was also reported to be nuclear-localized and its overexpression resulted in reduced peroxisome number (32). Notably, the genome of Arabidopsis contains one close homolog to BORI1, At4g14490, which we named BORI2. Both BORI1 and 2 are characterized by an N-terminal FHA domain and a helical domain at the C-terminus (Fig. 1*C-D*). The forkhead-associated (FHA) domain is a small protein module shown to recognize different phospho-epitopes on proteins, with a preference for phosphothreonine. It has been identified in both prokaryotes and eukaryotes in a diverse range of proteins such as kinases, phosphatases, RNA- and DNA-binding proteins as well as metabolic enzymes (33–36). To get hints on the putative function of the BORIs, we performed a phylogenetic analysis of the FHA domain of BORI1 and 2 (see Material and Methods). We found that BORI1 and 2 originated by a duplication in the common ancestor of Brassicacaea (Fig. 1*D*), and that additional BORI orthologs can only be found in Archaeplastida amongst both Chlorophyta (green algae) and Streptophyta (land plants), but not Rhodophyta (red algae). BORI orthologs are specifically characterized by the presence of a conserved C-terminal helix (Fig. 1*D*). Using the presence of this domain, we could separate BORI orthologs from its closest paralogous FHA domain, found in the PP2C phosphatase KAPP (kinase associated protein phosphatase (37)), which resulted from a duplication in the common ancestor of Viridiplantae (composed of Chlorophyta and Streptophyta). KAPP interacts with various receptor kinases, and regulates local phosphorylation status of such receptors at the plasma membrane (38, 39). The closest outgroup to KAPP/BORI FHA domains contains deltaproteobacterial proteins, suggesting a potential lateral transfer of FHA proteins from these prokaryotic lineages to the ancestor of Viridiplantae. The function of these prokaryotic homologs however, is unclear.

To further test and validate the interaction of both BORIs with BORR, we generated plants producing GFP fusion proteins of BORI1 and 2 (*PRO_BORI1_:BORI1:GFP, PRO_BORI2_:BORI2:GFP*, see below), and crossed them with plants expressing *PRO_BORR_:BORR:RFP*. As a control, we combined the previously generated *PRO_BORR_:BORR:RFP* plants with plants producing GFP alone (*PRO_35S_:GFP*) (21, 40). After immunoprecipitation with GFP-Trap beads using protein extracts from seedlings, we detected complex formation between BORR:RFP and BORI1:GFP as well as BORR:RFP and BORI2:GFP but not between BORR:RFP and GFP alone (Fig. 2*A*).

**Figure 2.**
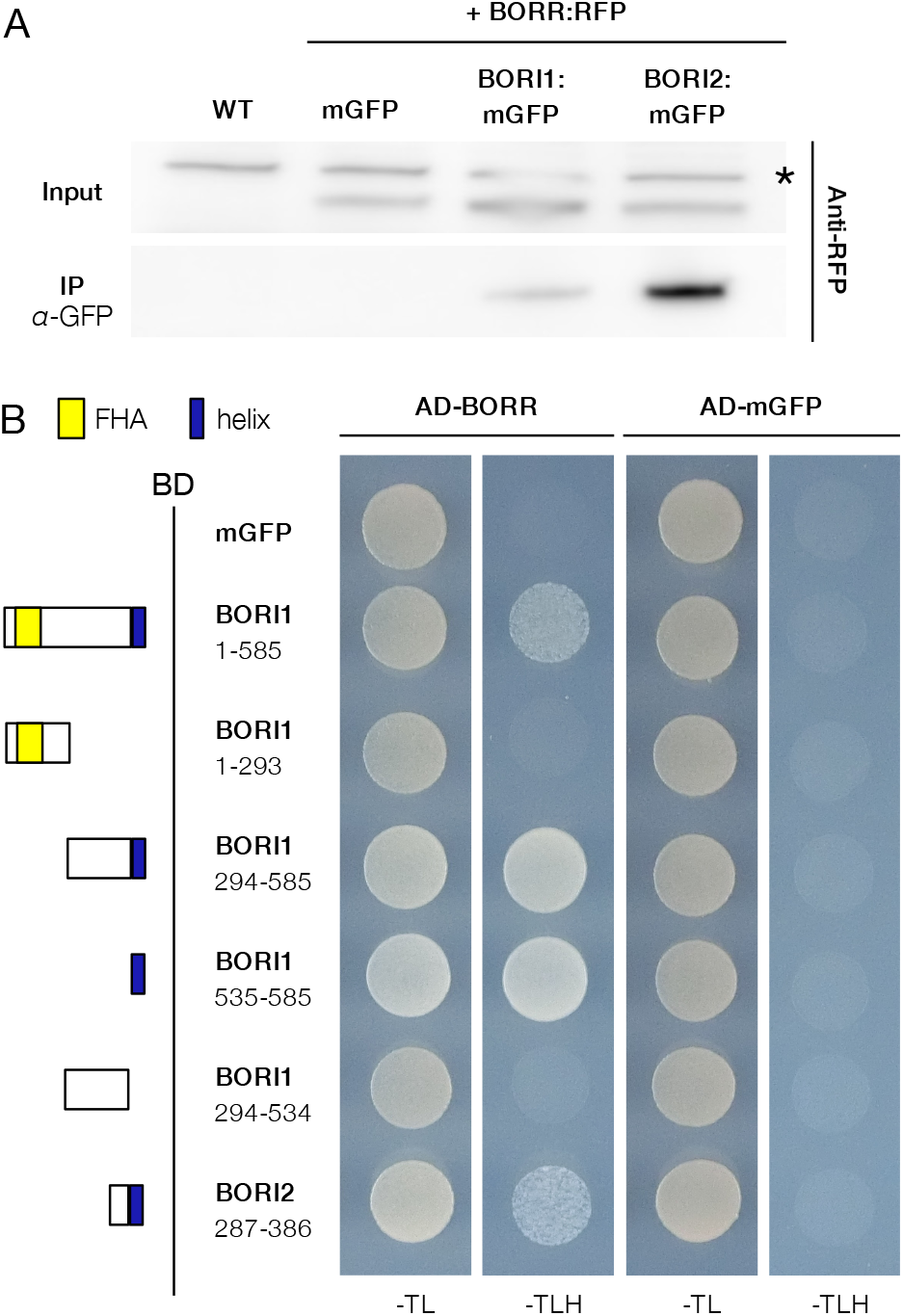
Interaction between BORIs and BORR. (**A**) Co-immunoprecipitation of BORIs and BORR from stable transgenic plants. 7-day-old Arabidopsis seedlings expressing BORR:RFP and BORI1:mGFP or BORR:RFP and BORI2:mGFP were used for IP with an anti-GFP antibody. Both input and IP fractions were subjected to immunoblotting with an anti-RFP antibody. Seedlings expressing both *BORR:RFP* and *mGFP* as well as WT seedlings were used as negative controls. The asterisk indicates a nonspecific band. (**B**) Identification of the interaction domain between BORIs and BORR by yeast two-hybrid assay. Each strain was spotted on SD plates without Trp and Leu (-TL; control media) or without Trp, Leu, and His (-TLH; selection media) and photographed after incubation at 30°C for 2 days. AD, GAL4-activation domain. BD, GAL4-DNA binding domain. mGFP was used as a negative control.

To test for direct interaction and to subsequently map the interaction domains between BORR and the BORIs, we performed yeast two-hybrid assays. Deletion analyses revealed that the conserved C-terminal helix of the BORIs is necessary for the contact with BORR (Fig. 2*B*). Since this domain configuration is reminiscent of Survivin, which also interacts with Borealin via its C-terminal helix in the context of the three helical bundle formed with INCENP (6), we speculated that the BORIs might be bona fide homologs of Survivin in plants by virtue of their C-terminal domains.

### The FHA domain of BORIs specifically binds phosphorylated histone H3 threonine 3

In case of functional conservation, we hypothesized that BORI, like Survivin, should be able to bind to phosphorylated H3T3 presumably via its N-terminal FHA domain. To test this, we first performed co-IP assays using protein extracts of transgenic Arabidopsis seedlings expressing *PRO_BORI1_:BORI1:GFP* or *PRO_BORI2_:BORI2:GFP*. Wild-type and *PRO_35S_:GFP*-expressing plants were used as negative controls. After immunoprecipitation with GFP-Trap beads, we successfully detected Histone H3 in BORI1:GFP and BORI2:GFP samples but not in the wild-type and GFP-alone samples (Fig. 3*A*). Next, we performed histone H3 peptide-binding assays to address whether BORIs can interact with phosphorylated histone H3 tails. Synthesized histone-H3 tails with or without a phosphate group at amino acid T3 and/or T11 were conjugated with biotin and incubated with the GST-fused FHA domain of BORI1 or BORI2. Bound proteins were retrieved using streptavidin-coupled magnetic beads and detected by western blotting. As shown in Figure 3*B*, both FHA domains weakly bound to non-phosphorylated H3 peptides. Notably, the affinities to H3 peptides were strongly enhanced by H3T3^ph^ but not H3T11^ph^, indicating that the FHA domain of BORIs can specifically recognize the phosphorylation status of histone H3 at threonine 3.

**Figure 3.**
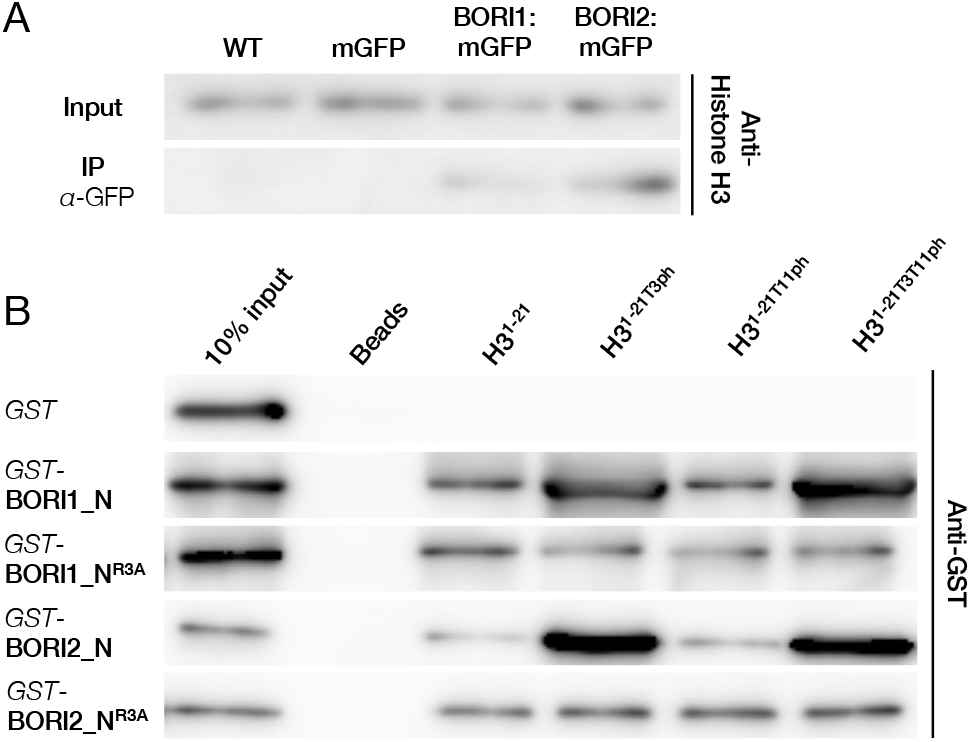
The FHA domain of BORIs directly binds to H3T3^ph^. (**A**) Co-immunoprecipitation of BORIs and Histone H3 from stable transgenic plants. 7-day-old Arabidopsis seedlings expressing BORI1:mGFP or BORI2:mGFP were used for IP with an anti-GFP antibody. Both input and IP fractions were subjected to immunoblotting with an anti-Histone H3 antibody. Seedlings expressing mGFP and WT seedlings were used as negative controls. (**B**) Peptide-binding assay. All peptides were biotinylated at the C-terminus and were preincubated with streptavidin-coated beads before addition of FHA domains. Protein binding was subjected to immunoblotting with an anti-GFP antibody. GST alone was used as a negative control.

According to different structures of FHA domains bound to phosphopeptides, arginine residues in the loops between the ß-sheets of the FHA domain are often involved in direct contact with the phosphate residue of the phospohopeptide, including those found in the FHA domain of KAPP (41–44). Using the Alphafold2 3D predicted structures of BORI1 and 2 (*SI Appendix*, Dataset S2), we identified three arginine residues present in loops that are potentially facing the phosphorylated histone H3 (Fig. 1*C-D*). We substituted all three arginine residues in these loops with alanine (Fig. 1*C*) with the aim to disturb H3T3^ph^ binding. When tested *in vitro*, both mutated FHA domains (R3A mutants) indeed showed drastically reduced H3T3^ph^-binding affinities (Fig. 3*B*).

### BORIs are required for proper chromosome segregation and cell division

For a functional analysis, we isolated a T-DNA insertion mutant of *BORI1* (*bori1-1*) that did not express full-length *BORI1* transcript (*SI Appendix*, Fig. S1*A-C*). Since T-DNA insertion mutants for *BORI2* were not available, we generated a mutant by CRISPR/Cas9. The resulting *bori2-1* allele has an 8-bp deletion that creates a premature stop codon (*SI Appendix*, Fig. S1*A*). Since both single mutants showed no obvious mutant phenotypes, we combined *bori1* with *bori2* to overcome a possible functional redundancy. No difference in growth and fertility in comparison to the wildtype were detected in mutants which are homozygous for one and heterozygous for the other *bori* mutant allele (*SI Appendix*, Fig. S2*A-C*). However, double homozygous mutants could not be recovered among more than 380 seedlings in the progeny of *bori1^+/-^ bori2^-/-^* and *bori1^-/-^ bori2^+/-^* plants. Consistently, we observed aborted seeds and undeveloped ovules in siliques of *bori1^+/-^ bori2^-/-^* and *bori1^-/-^ bori2^+/-^* (Fig. 4*A* and *B*).

**Figure 4.**
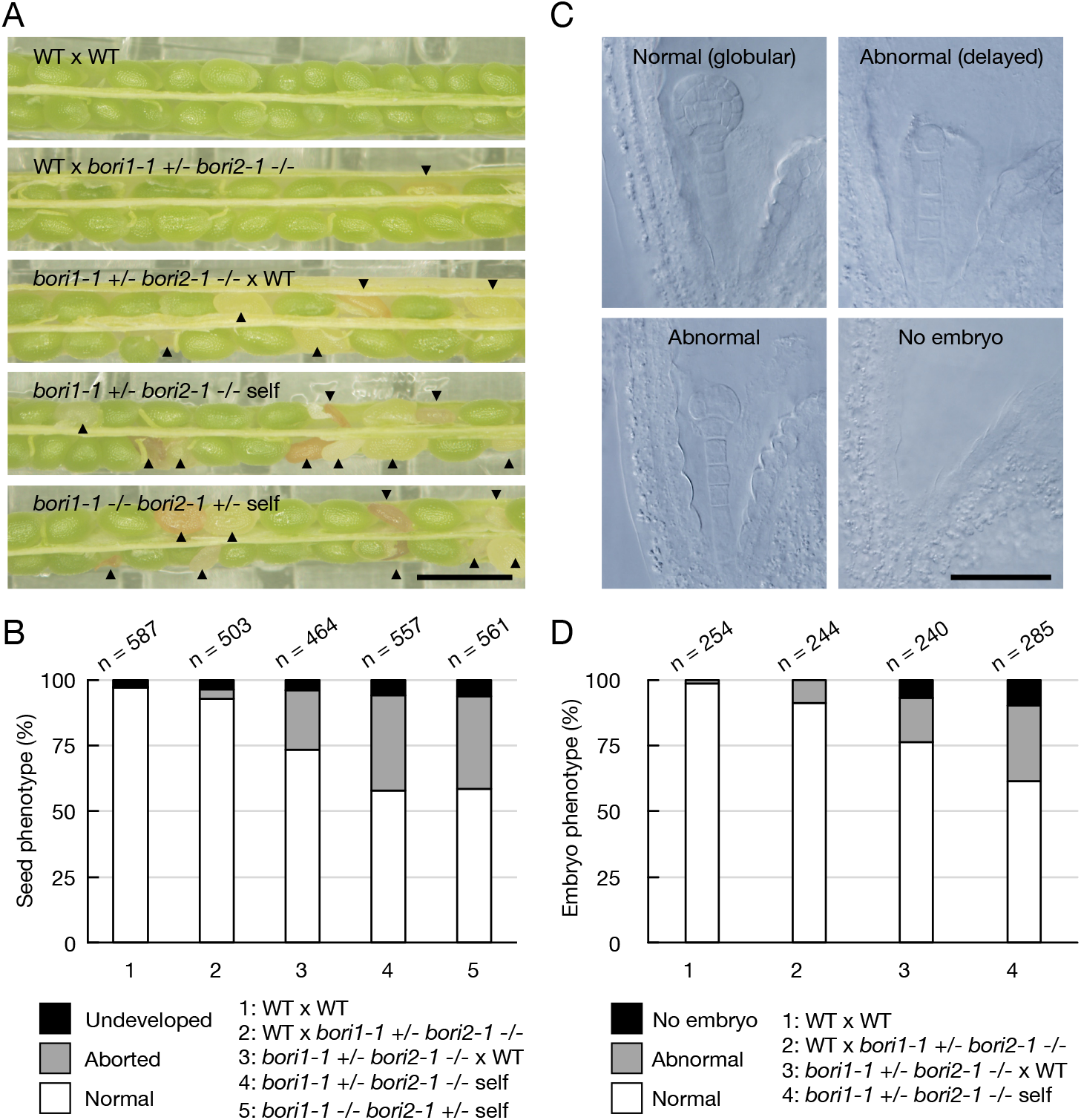
Mutants in *BORI* exhibit defects in embryo development. (**A**) Developing seeds in a silique resulting from reciprocal crosses between *bori* mutants and the WT. Arrowheads indicate aborted seeds. Scale bar, 1 cm. (**B**) Frequency of seed phenotypes shown in each cross. (**C**) Embryo phenotypes observed in *bori* mutants. Whole-mount clearing was conducted 4 days after the pollination. Scale bar, 100 μm. (*D*) Frequency of embryo phenotypes shown in each cross.

To confirm that the lethal phenotype was due to the mutated *BORI* genes, we carried out complementation tests using the two *BORI* genomic fragments, each fused with *GFP* (*PRO_BORI1_:BORI1:GFP* and *PRO_BORI2_:BORI2:GFP*, see above). Either construct could fully complement the lethal phenotype of the double homozygous mutants, indicating that *BORI* function is essential in plants, similarly to the previously analyzed other two components of the CPC (*SI Appendix*, Fig. S2*A-C*) (19, 21).

Since we observed more than 25% aborted and/or undeveloped seeds as well as more than 25% seeds with an abnormal embryo or no embryo (Fig. 4*B* and *D*), we performed a reciprocal cross between *bori1^+/-^ bori2^-/-^* and wild-type plants to examine the transmission efficiency of the *bori1* allele as a proxy for developmental defects of the mutant gametophytes. When we used *bori1^+/-^ bori2^-/-^* as the male plant, the transmission efficiency was 94.2% (*n* = 200). Conversely, when we used *bori1^+/-^ bori2^-/-^* as the female plant, the transmission efficiency was reduced to 38.9% (*n* = 200), suggesting that BORIs have an important role in the function and/or development of the female gametophyte. However, when analyzing the mature siliques, we rarely observed unfertilized ovules, indicating that this reduction of the transmission efficiency manifested only after fertilization during embryo development (Fig. 4*A* and *B*). Consistently, when *bori1^+/-^ bori2^-/-^* was used as the female parent and pollinated with wild-type pollen, we frequently found abnormal embryo development (Fig. 4*C* and *D*).

To study the *BORI* function after embryogenesis, we constructed an artificial microRNA against *BORI2* (*amiBORI2*), and transformed it into *bori1^-/-^* mutants. Most of the transformants (15 of 18) exhibited a dwarf phenotype. We selected two transformants that showed approximately 34% and 66% reduction of *BORI2* transcript levels for further analysis (*amiBORI2#1^bori1^* and *amiBORI2#2 ^bori1^*) (*SI Appendix*, Fig. S1*D*). Both mutants exhibited a dose-dependent reduction in leaf area and curling of leaves (*SI Appendix*, Fig. S2*A*). As observed in *BORR* knockdown mutants, the reduction of *BORI* transcript levels also resulted in plants displaying the so-called *bonsai* phenotype with compact inflorescences at flowering stage (*SI Appendix*, Fig. S2*B* and *C*). To further test whether compromised CPC function gives rise to a *bonsai* phenotype, we also generated plants expressing a microRNA against AUR3 (*SI Appendix*, Fig. S1*E*). Indeed, *AUR3* knockdown plants revealed the same phenotype as *amiBORR* and *amiBORI* mutant plants (*SI Appendix*, Fig. S2*A-C*).

The growth of primary roots was also compromised by the knockdown of *BORI2* in *bori1* mutants (Fig. 5*A* and *B*). We observed that many cells died in the root meristems of both *amiBORI2#1^bori1^* and *amiBORI2#2^bori1^* plants (Fig. 5*C* and *D*). In addition, both mutants produced small root meristems with aberrant cell division (Fig. 5*C, E* and *F*). We have previously shown that impaired CPC activity in plants causes chromosome segregation defects, for instance lagging chromosomes in anaphase, consistent with compromised CPC activity in animals and yeast (21). To monitor possible mitotic defects, we crossed the *amiBORI2#1^bori1^* and *amiBORI2#2^bori1^* mutants with a previously generated transgenic line expressing both a microtubule (*RFP:TUA5*) and a centromere (*GFP:CENH3*) marker (40). Analysis of the resulting plants revealed that both *amiBORI2#1^bori1^* and *amiBORI2#2^bori1^* mutants also have lagging chromosomes and the frequency of these segregation defects increased with the level of *BORI* transcript reduction (Fig. 5*G* and *H*). Thus, loss of BORI function results in aneuploidy and likely in the further course of development, secondary developmental defects and cell death. This finding also could explain the severe embryonic versus rather mild gametophytic defects described above since aneuploidy arisen during gametophyte development might cause embryo abortion only after a few divisions of the zygote.

**Figure 5.**
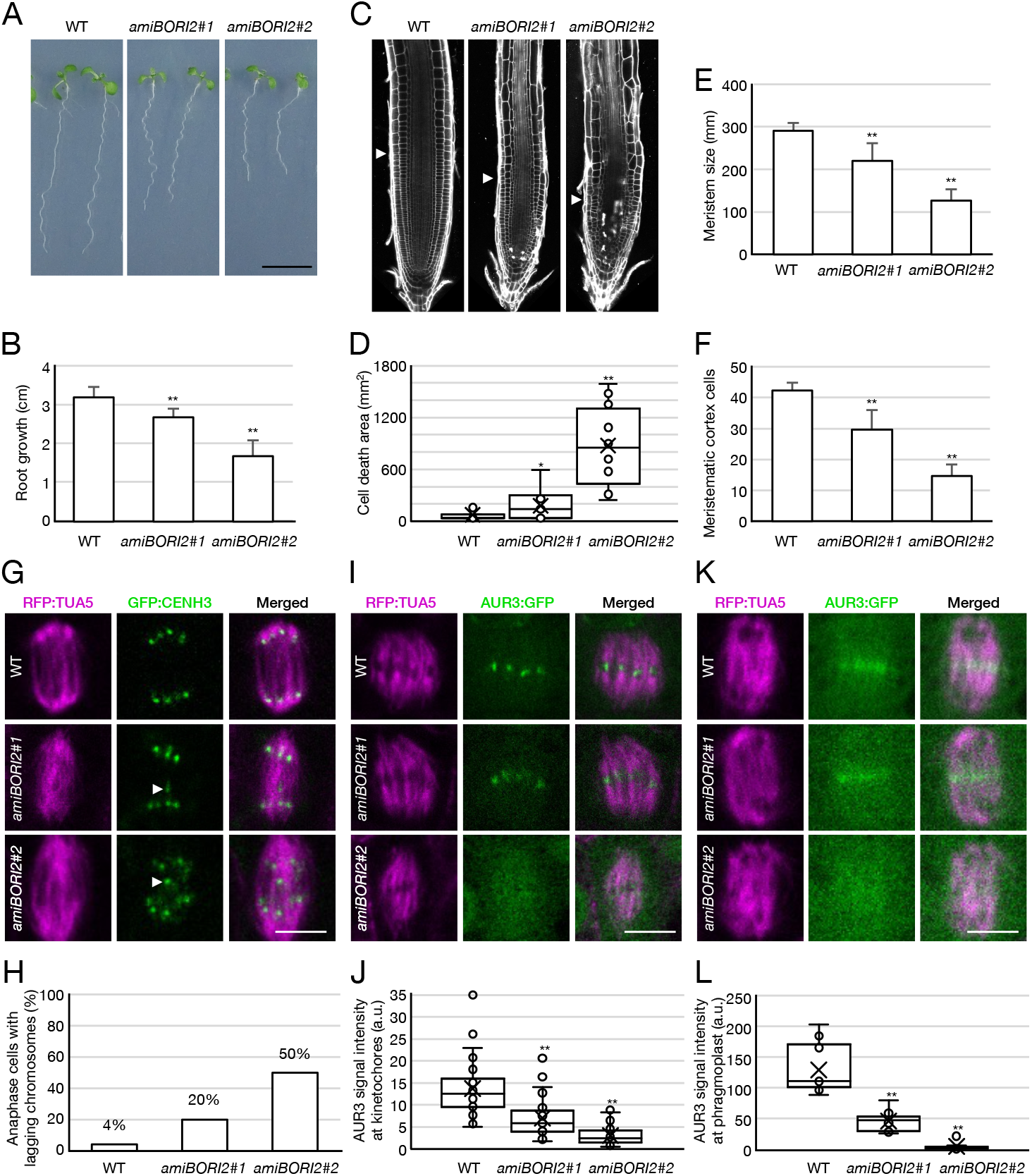
Phenotypic analysis of amiRNA-mediated *BORI* knockdown plants. (**A**) 7-day-old WT and *BORI* knockdown seedlings. Scale bar, 1 cm. (**B**) Root length of 7-day-old WT and *BORI* knockdown seedlings. Graph bars represent means ± SD. Asterisks indicate significant difference between the WT and *BORI* knockdown seedlings tested by Student’s t-test (\*\**P* < 0.001, *n* = 30). (**C**) Confocal images of 7-day-old WT and *BORI* knockdown roots stained with 20 μg/ml propidium iodide to visualize cell walls and dead cells. Arrowheads indicate the boundary between the division region and the elongation region of the root. Scale bar, 100 μm. (**D**) Cell death area in *C*. Asterisks indicate significant difference between WT and *BORI* knockdown seedlings tested by Student’s t-test (\**P* < 0.01, \*\**P* < 0.001, *n* = 20). (**E**) Meristem size in *C*. Meristem size was measured from quiescent center to the first elongated cell in the cortical cell file. Asterisks indicate significant difference between WT and *BORI* knockdown seedlings tested by Student’s t-test (\*\**P* < 0.001, *n* = 20). (**F**) Number of meristematic cortex cells in *C*. Asterisks indicate significant difference between WT and *BORI* knockdown seedlings tested by Student’s t-test (\*\**P* < 0.001, *n* = 20). (**G**) Representative images of normally distributed and lagging chromosomes in 5-day-old WT and *BORI* knockdown roots. Microtubules and centromeres were visualized by RFP:TUA5 and GFP:CENH3, respectively. Arrowheads indicate lagging chromosomes. Scale bar, 5 μm. (**H**) Frequency of lagging chromosomes in anaphase cells in *G*. *n* = 50. (**I**) Representative images of AUR3 accumulation levels at metaphase centromeres in 5-day-old WT and *BORI* knockdown roots. Microtubules and AUR3 were visualized by RFP:TUA5 and AUR3:GFP, respectively. Scale bar, 5 μm. (**J**) AUR3 signal intensity in *I*. AUR3:GFP signals at metaphase centromeres were measured. Asterisks indicate significant difference between WT and *BORI* knockdown roots tested by Student’s t-test (\*\**P* < 0.001, *n* = 50). (**K**) Representative images of AUR3 accumulation levels at the middle part of the phragmoplast in 5-day-old WT and *BORI* knockdown roots. Microtubules and AUR3 were visualized by RFP:TUA5 and AUR3:GFP, respectively. Scale bar, 5 μm. (**L**) AUR3 signal intensity in *K*. AUR3:GFP signals at the middle part of the phragmoplast were measured. Asterisks indicate significant difference between WT and *BORI* knockdown roots tested by Student’s t-test (\*\**P* < 0.001, *n* = 10).

To address whether the *BORI* loss-of-function phenotype was due to mislocalization of AUR3, we investigated the AUR3 accumulation at centromeres and phragmoplast by introducing a *PRO_AUR3_:AUR3:GFP* reporter (21) into *amiBORI2^bori1^* plants. Indeed, abundance of AUR3 at centromeres and phragmoplast was drastically reduced in both *amiBORI2#1^bori^* and *amiBORI2#2^bori1^* mutants, indicating that BORIs are required for proper localization of the CPC complex (Fig. 5*I-L*).

### BORIs are needed for proper targeting of the CPC to chromatin *in vivo*

Transient expression of the *BORI1* cDNA fused to the ORF of GFP under the control of the constitutive *Cauliflower mosaic virus promoter 35S* was previously found to result in homogeneous accumulation of the fusion protein in the nucleoplasm of mature tobacco leaves (32). To investigate the detailed subcellular localization of BORIs during cell proliferation, we combined each of our genomic *BORI:GFP* reporters with a *RFP:TUA5* marker line, and analyzed their co-expression pattern in Arabidopsis root tips. BORI1:GFP and BORI2:GFP showed the same localization pattern throughout the cell cycle (Fig. 6*A*, *SI Appendix*, Fig. S3*A*, and Movie S1 and 2) and thus, will be described in the following as BORI:GFP. In interphase, BORI:GFP localized to the nucleoplasm and accumulated in nuclear foci. Co-localization analyses with *RFP:CENH3* revealed that these foci are centromeres. Since CENH3 marks the kinetochore-proximal centromeres, we further concluded that BORI:GFP resides at inner centromeres similar to the other CPC components (*SI Appendix*, Fig. S3*C*) (21).

**Figure 6.**
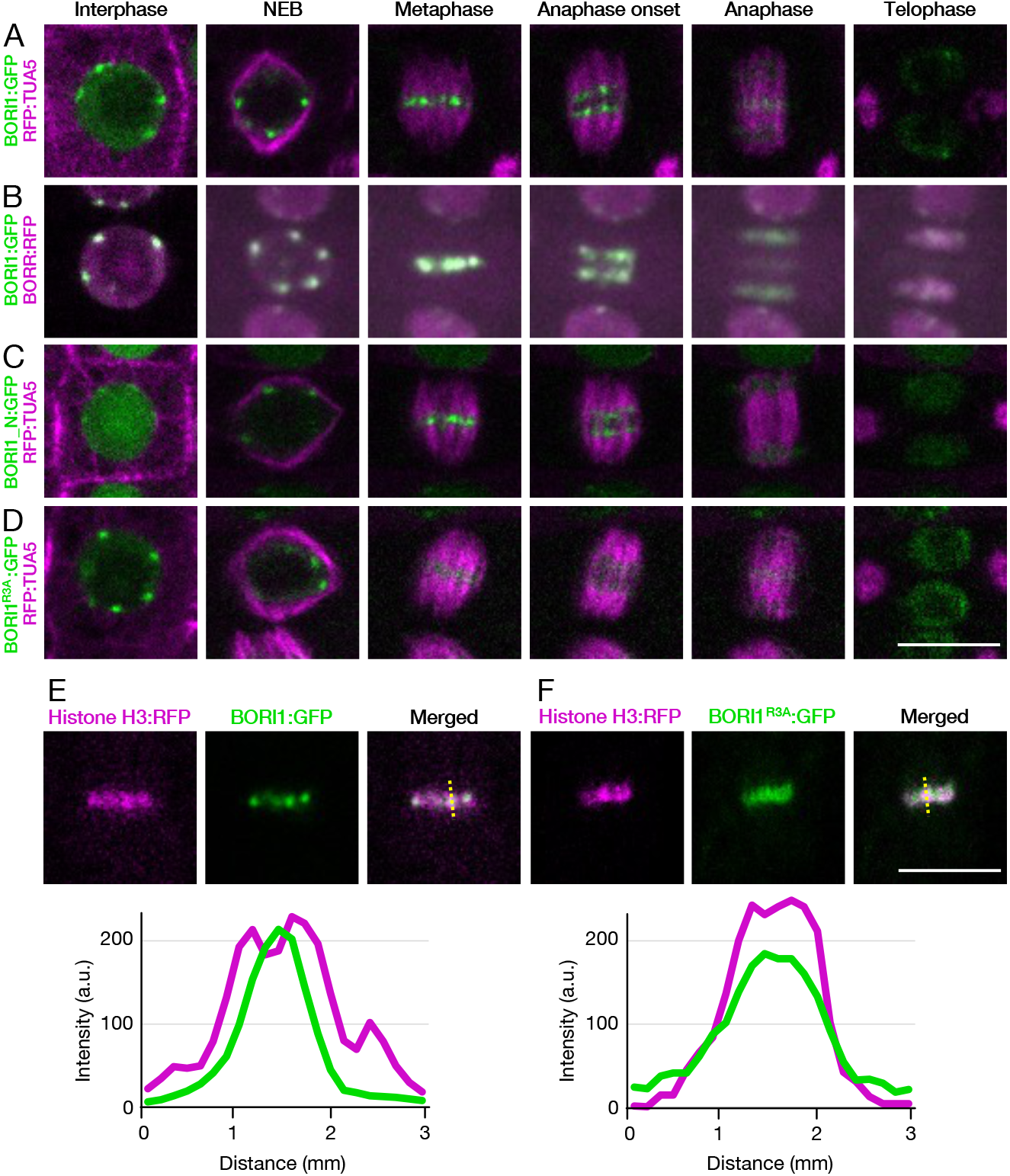
Subcellular localization of BORI1 during the cell cycle. (**A**) Subcellular localization of BORI1:GFP during the cell cycle. Microtubule structures were visualized by RFP:TUA5. (**B**) Colocalization of BORI1:GFP and BORR:RFP. (**C and D**) Subcellular localization of BORI1_N:GFP (**C**) or BORI1^R3A^:GFP (**D**) during the cell cycle. Microtubule structures were visualized by RFP:TUA5. For live imaging, root tips of 5-day-old seedlings were used. Scale bar, 10 μm. (**E and F**) Colocalization of Histone H3:RFP and BORI1:GFP (**E**) or BORI1^R3A^:GFP (**F**) in metaphase cells. The yellow dotted line indicates the positions where the line profiles were obtained. Scale bar, 10 μm.

In mitotic cells, the BORI:GFP signal rapidly concentrated at the centromeres starting just after nuclear envelope breakdown until metaphase. After anaphase onset, BORI:GFP changed its localization from centromeres to the center of phragmoplasts, and, after completion of cell division, BORI:GFP re-accumulated in the nucleus. The localization pattern of BORI1:GFP completely overlapped with that of BORR:RFP throughout the cell cycle (Fig. 6*B* and Movie S3).

To understand the spatial regulation of BORI during the cell cycle, we generated a BORI1:GFP reporter driven by its own promoter but without the BORR-interacting helix at the C-terminus of the protein (amino acid 1-293, called *PRO_BORI1_:BORI1_N:GFP* in the following) BORI1_N:GFP accumulated at centromeres from prophase to metaphase similar to the full length BORI1 fusion with GFP. However, BORI1_N:GFP signals were only detected on chromosomes and not at the phragmoplast in late anaphase (Fig. 6*C* and Movie S4). These results indicate that the interaction of BORIs with BORR is necessary for the targeting of BORIs to phragmoplasts but not to centromeres.

In other organisms, Survivin concentration at inner centromeres relies on the phosphorylation of histone H3 at threonine 3 (H3T3^ph^) catalyzed by Haspin. To check whether the H3T3^ph^ mark is also required for proper BORI localization to centromeres, *BORI1:GFP* expressing plants were treated with 5-Iodotubercidin (5-ITu), a commonly used Haspin inhibitor (23, 45). Although we could still detect BORI1:GFP at centromeres after treatment, the GFP signal in metaphase was much more diffuse (*SI Appendix*, Fig. S3*D*). Notably, we could reproduce the same localization defect by introducing the R3A substitution into *BORI:GFP* (*BORI^R3A^:GFP*) that reduces the binding affinity to H3T3^ph^ (see above) (Fig. 6*D*, *SI Appendix*, Fig. S3*B*, and Movie S5 and 6). To analyze the localization of BORI^R3A^:GFP in detail, we created transgenic plants that expressed a Histone H3 marker (Histone H3:RFP) together with BORI:GFP or BORI^R3A^:GFP. Whereas BORI:GFP only colocalized with Histone H3:RFP at the inner region of the centromere, BORI^R3A^:GFP localized to the entire Histone H3:RFP-marked region (Fig. 6*E* and *F*). These results demonstrate that the binding of BORIs to H3T3^ph^, similar to Survivin in animals and yeast, is crucial for their accumulation at the inner centromere.

To address the functional relevance of the precisely targeted centromere localization of BORIs, we expressed the *BORI1^R3A^:GFP* construct in *bori1 bori2* double homozygous mutants. While the *BORI1^R3A^:GFP* construct could complement the lethal phenotype of the double homozygous mutants, the resulting plants exhibited a wide range of developmental defects (Fig. 7*A* and *B* and *SI Appendix*, Fig. S2*D-F*). Foremost, these plants displayed severe growth defects and, as observed in *BORI* knockdown plants, the *BORI1^R3A^:GFP* expressing plants also contained many dying cells in their root meristems possibly caused by aneuploidy (Fig. *7C* and *D*). Indeed, we observed chromosomal variations in *BORI1^R3A^:GFP-expressing* root cells that contained between 8 and 11 chromosomes in contrast to the invariable 10 chromosomes found in the wildtype (Fig. 7*E* and *F*, and Movie S7-9). These results demonstrate that proper centromere localization of BORIs, which is mediated by their recognition of H3T3^ph^, is required for genome stability.

**Figure 7.**
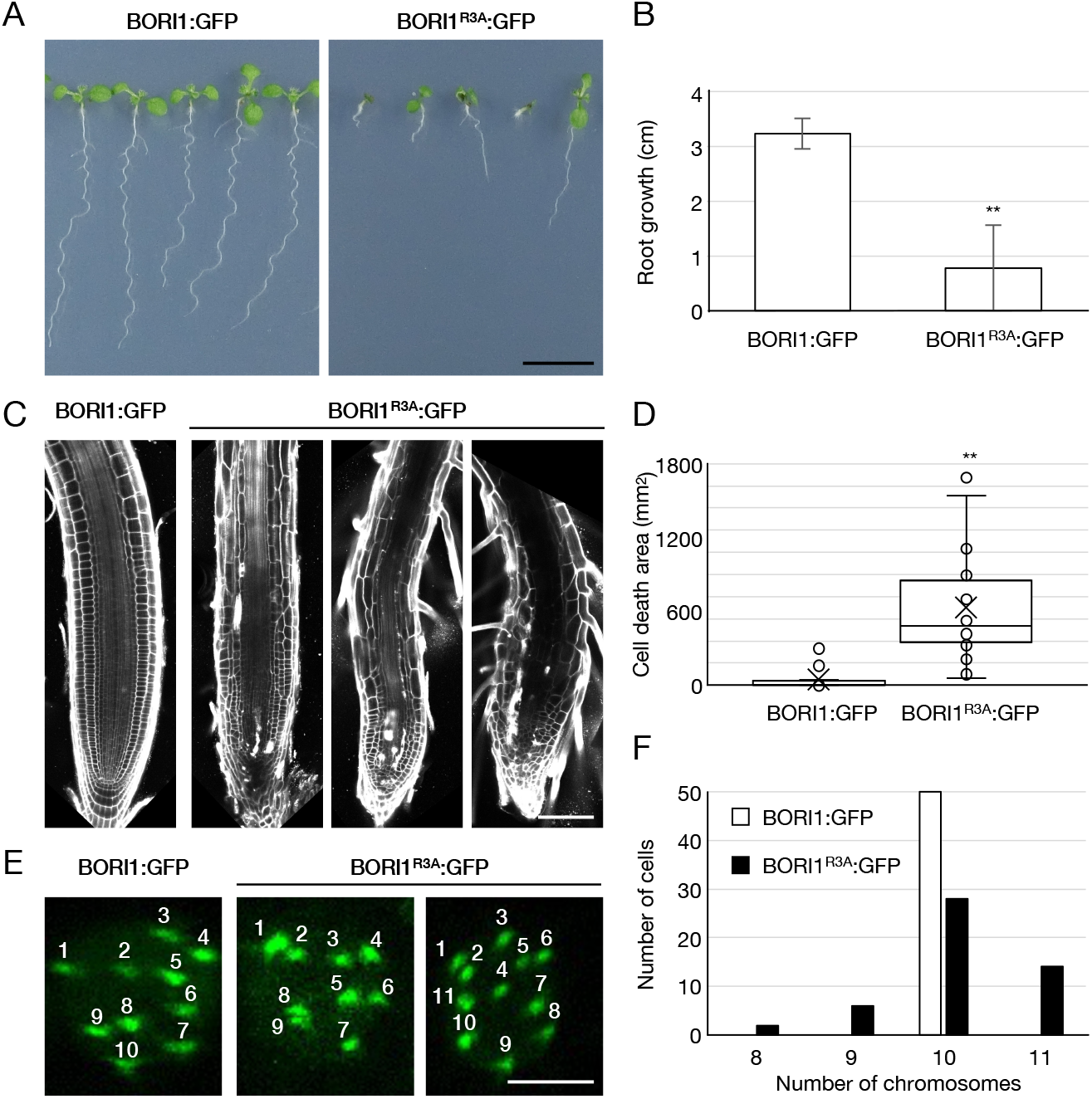
Proper centromere localization of BORIs is required for genome stability. (**A**) 7-day-old transgenic lines expressing *BORI1:GFP* or *BORI1^R3A^:GFP* in *bori1 bori2*. Scale bar, 1 cm. (**B**) Root length of 7-day-old transgenic seedlings. Graph bars represent means ± SD. Asterisks indicate significant difference tested by Student’s t-test (\*\**P* < 0.001, *n* = 30). (**C**) Confocal images of 7-day-old *BORI1:GFP* and *BORI1^R3A^:GFP* seedling roots stained with 20 μg/ml propidium iodide to visualize cell walls and dead cells. Scale bar, 100 μm. (**D**) Cell death area in *C*. Asterisks indicate significant difference tested by Student’s t-test (\*\**P* < 0.001, *n* = 20). (**E**) GFP signals in interphase cells of *BORI1:GFP* and *BORI1^R3A^:GFP* seedling roots. Scale bar, 5 μm. (**F**) Number of chromosomes in interphase cells of *BORI1:GFP* and *BORI1^R3A^:GFP* seedling roots (*n* = 50).

### The defining feature of the Survivin/BORI gene family is a coiled-coil forming helix

Since we found that the BORIs behave analogous to Survivin in animals and fungi with respect to their function in the CPC, and that both Survivin and BORIs harbor a C-terminal conserved helix, we wondered whether these helices would show significant sequence similarity indicating a common evolutionary history. We therefore constructed Hidden Markov Models (HMMs), using the Arabidopsis BORI C-terminal helices as a template. Following iterative searches in large eukaryotic sequence databases using our trained HMM, we identified bona fide animal and fungal Survivin homologs, while known unrelated helices were never picked up under the filtering conditions used (for criteria see Material and Methods). This strategy also worked in a reciprocal fashion, i.e., when iterative similarity searches were initiated with the helix of animal Survivin homologs, we were able to identify plant BORI homologs. Such reciprocal similarity connections strongly suggested that Survivin and BORI are bona fide orthologs, belong to the same gene family, and that the BIR/FHA domains should be considered accessory domains. Excitingly, using our helix-specific HMM-based strategy, we could identify Survivin/BORI orthologs in all eukaryotic supergroups in which no Survivin-like protein was previously found (Fig. 8*A-B*), as earlier searches concentrated on the more distinct BIR domain of animal and fungal Survivins (20). We detected orthologs with four different domain topologies in different eukaryotic clades (Fig. 8*A-D*): (I) Fungi and Metazoa display the known Survivin structure of 1 or 2 N-terminal BIR domains and a C-terminal helix, (II) Viridiplantae harbor an N-terminal FHA domain and a C-terminal helix, (III) Stramenopila and Haptista have a reversed topology with an FHA domain at the C-terminus following the conserved helix, (IV) finally, Amoebozoa, Rhodophyta, Discoba, Cryptista and Metamonada contain relatively short homologs, which only comprise the helix but no additional recognisable domains, as the phosphate-binding FHA or BIR domains. To determine the evolutionary history of the FHA domain of BORI-like orthologs found in Stramenopila and Haptista, relative to those found in Viridiplantae BORIs, we performed a phylogenetic analysis (see Material and Methods). We found that both FHA sub-types present amongst BORI-like orthologs are closely related to different FHA domains found in Deltaproteobacteria (Fig. 8*C*). Such phylogenetic relationships indicate an ancient lateral transfer of prokaryotic FHA domains to at least two ancient eukaryotic ancestral lineages, suggesting a distinct, but ancient prokaryotic evolutionary origin for the FHA domain found in BORI-like orthologs of Viridiplantae as well as Stramenopila and Haptista. Our phylogenetic analyses and sensitive sequence searches point to an evolutionary scenario for the Survivin/BORI gene family in which the ancestral version present in the last eukaryotic common ancestor (LECA) only consisted of a helix contributing to the triple helix structure of the CPC scaffold. The acquisition of a phosphate-binding domain occurred independently in at least three different clades resulting in molecular convergence between Survivin in Fungi and Metazoa (+N-terminal BIR), BORI-like orthologs in Viridiplantae (+N-terminal FHA), and Stramenopila and Haptista (+C-terminal FHA) with respect to their capacity to bind phosphorylated H3T3 (Fig. 8*D*).

**Figure 8.**
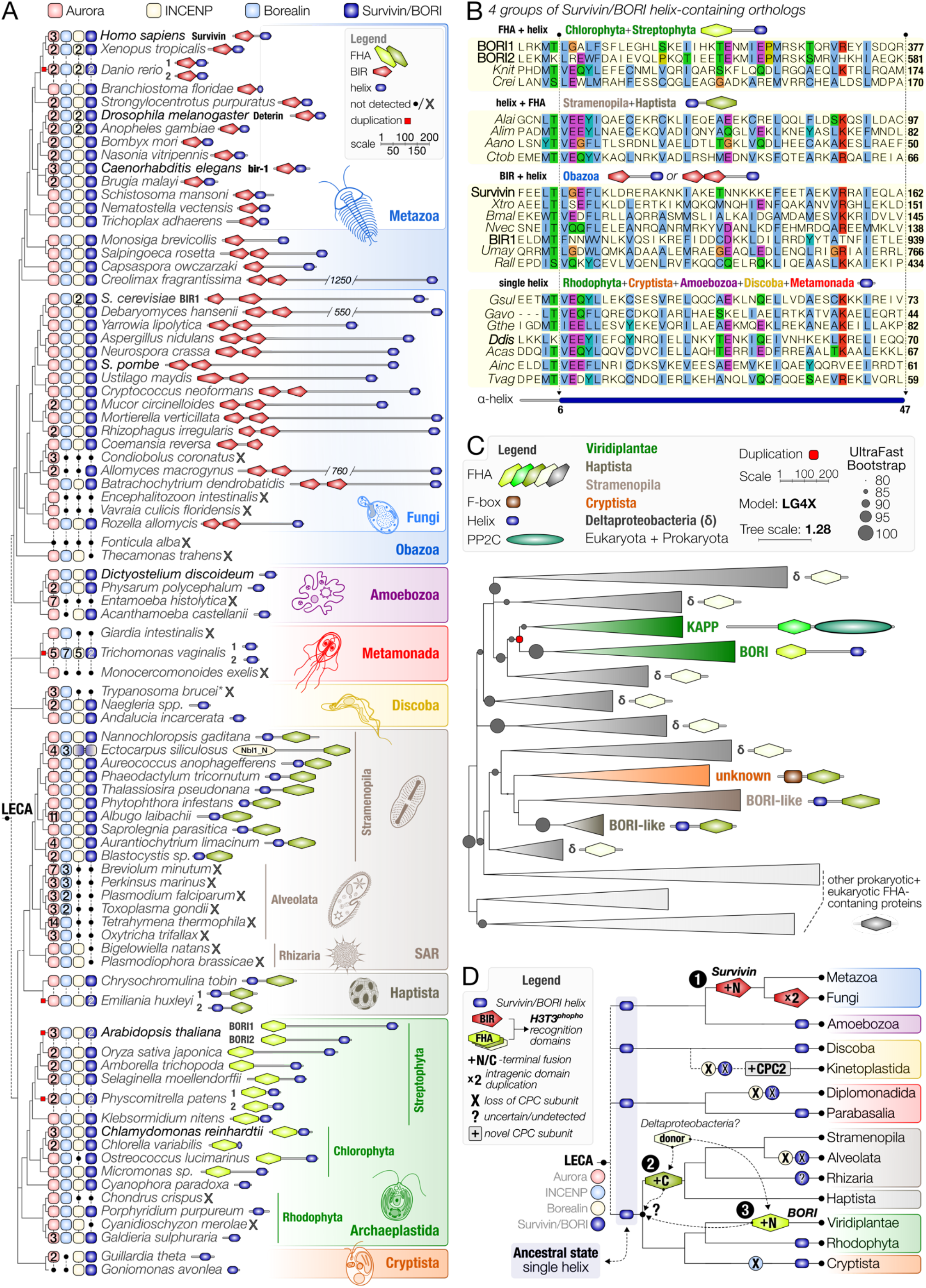
A conserved helix and the recurrent acquisition of a phosphate-binding domain characterize the divergent Survivin/*BORI* gene family. (**A**) Presence-absence matrix of the four subunits of the CPC: Aurora kinase, INCENP, Borealin, and Survivin/BORI in a wide variety of eukaryotes, with representatives of all supergroups (different colors) found across the eukaryotic tree of life (according to Burki *et al*. 2019 (71)). Dashed lines indicate uncertain relationships amongst supergroups. LECA refers to the hypothetical position of the last eukaryotic common ancestor. Numbers indicate the paralogs present in a specific clade. Domain topologies for the Survivin/BORI orthologs are shown on the right, with the presence of a conserved helix (blue) as the defining feature of the Survivin/BORI gene family. Different colors for the FHA domains indicate a unique evolutionary origin of those present in BORI orthologs in Archaeplastida as well as SAR and Haptista. (**B**) Multiple alignment of a conserved helix found in four sub-types of Survivin/BORI orthologs found amongst eukaryotes. Four letter abbreviations refer to species that can be found in panel A (a combination of the first letter of the genus name and the first three letters of the species name). A small vertical space in the alignment (e.g. between Aano and Ctob, indicate that those species are part of a different supergroup: SAR and Haptista, respectively). Numbers on the right indicate the position of the most C-terminal residue of a 47-residue-long conserved helix. (**C**) Collapsed and unrooted maximum-likelihood phylogenetic tree of eukaryotic and prokaryotic FHA domains most similar to BORI, and Survivin/BORI-like proteins found amongst Archaeplastida, SAR, and Haptista. Representative domains for each collapsed clade are projected onto the phylogenetic tree (see legend). Branch lengths are scaled and indicate the number of substitutions per site (see scale bar). Circles indicate bootstrap support (1000x replicates, only higher than 80% support shown). Colors indicate the different evolutionary histories of various domains (e.g. FHA of KAPPs and BORIs). For phylogenetic analysis details, see *SI Appendix*, Dataset S4. (**D**) Evolutionary scenario of the Survivin/*BORI* gene family. A helix-only Survivin/*BORI* gene was present in LECA and independently fused to a phosphate-binding domain at least three different times during eukaryotic evolution, in the ancestors of Fungi and Metazoa (1:BIR), SAR and Haptista (2:C-terminal FHA), and Viridiplantae (3:N-terminal FHA). The FHA domains were likely vertically transferred from Deltaproteobacteria, Colors are similar to panel A and C.

## Discussion

A dynamic subcellular localization of the CPC is crucial for its functions in the control of chromosome segregation and cytokinesis. To fulfill these tasks, INCENP, Borealin, and Survivin form a scaffold that guides the catalytic subunit Aurora kinase to its proper locations (Fig. 8). In animals and yeast, one of the scaffolding proteins, Survivin, has two indispensable functions for the CPC. First, Survivin interacts with the other scaffold proteins through its C-terminal alpha helix by engaging in a three-helical bundle, tying these three proteins together. Second, it recognizes the phosphorylation status of Histone H3 tail at threonine 3, a function provided by the BIR domain of Survivin in animals and yeast. However, no BIR-domain containing proteins exist in the plant lineage and only a distantly related BLD-domain has been described in putative IAP paralogs, one of which is involved in DNA demethylation (27, 46). Therefore, Survivins, as part of the CPC, were speculated to be non-existent in plants (47).

Here, we have identified BORI1 and 2, two novel BORR-interacting proteins, that execute the Survivin function in the CPC of Arabidopsis in a redundant fashion. The BORIs belong to a Viridiplantae-specific protein family with members of variable size that consists of an N-terminal FHA domain and a conserved C-terminal helix with a propensity to form a coiled-coil (triple helix) structure. We demonstrate that the FHA domain of BORIs act as a H3T3^ph^ reader at inner centromeres, a function fulfilled by the BIR domain in animals and yeast. Although both domains are capable of binding H3T3^ph^, they are structurally very different. The FHA domain consists of a sandwich of ß-sheets while the BIR domain displays three short alpha-helices and is stabilized by a zinc molecule, tetrahedrally coordinated by one histidine and three cysteine residues. Thus, there is no evidence of a common evolutionary history of the two domains. In contrast, the C-terminus of the BORIs, which we showed here to interact with BORR, has residual sequence similarity with the helical domain of Survivin in animals and yeast as found by reciprocal HMM searches indicating a common ancestor.

Interestingly, the HMM searches based on the conserved helix also led to the discovery of Survivin/BORI-type proteins in additional phylogenetic groups which either display the helix as the only defined structure in a relatively short protein or the helix plus an FHA-domain at the C-terminal end of the protein (Fig. 8). Based on this, we propose an evolutionary scenario in which the LECA possessed a helix-only version of Survivin to tether the three structural components of the CPC together. Then, in a process of convergent evolution different H3T3^ph^-binding domains were recruited in the different branches of the eukaryotic domain, likely to optimize and focus CPC localization at the inner centromere. Since several mechanisms contribute to the defined centromere localization of the CPC during mitosis in animals, it is possible that these were added in a stepwise manner. It would be plausible, for example, that binding of the ancestral CPC to centromeres was originally mediated by Borealin interacting with nucleosomes in general and centromere-localized Shugoshin in particular, and that binding to H3T3^ph^ was only added later to optimize the localization or enhance the concentration of the CPC. Consistently, when we disturbed H3T3^ph^ binding by point-mutations in the FHA domain, BORI localization to chromosomes was weaker and not confined to the centromere region in comparison to the non-mutated version, but not completely lost. Notably, the mutant phenotype of *bori1 bori2* double mutants expressing the *BORI1^R3A^:GFP* construct is also less severe than for a null mutant in any of the CPC components and rather resembled the effect of expressing a BORI-RNAi construct. However, although H3T3^ph^ binding by Survivin-type proteins is not the only pathway for centromere localization, the fact that we observe convergent molecular evolution in animals and plants, and possibly also in Stramenopila and Haptista, where an FHA domain has been added at the C-terminus, suggests that binding to H3T3^ph^ is a very efficient way to concentrate the CPC at the centromeres.

Given that not all Survivin candidates identified by the HMM searches display a distinct phosphate-binding domain, it will be interesting to see whether these putative orthologs are still part of the CPC in their respective lineages and whether H3T3^ph^ binding is still somehow mediated by these helix-only Survivin/BORI candidates or taken over by another CPC component. Alternatively, CPC concentration at centromeres in these cases might only be dependent on Borealin and Shugoshin, or yet an unidentified pathway. Noteworthily, studies of Dictyostelium INCENP and Aurora kinase, for example, indicate a mitotic CPC localization to centromeres resembling the pattern found in animals and plants (48, 49). Yet, the detailed distribution within the centromeric region is unclear. While a Haspin homolog has been identified in Dictyostelium (50), a CPC-Haspin interdependency in this lineage is not clear yet. Thus, it will be interesting to follow up the function of the helix-only variant of Survivin present in this species.

Another question is raised by the observation that we were still not able to identify the Survivin/BORI-type scaffolding component of the CPC for some clades while other components are present (e.g., Alveolata and Rhizaria in Fig. 8). Is the CPC assembled differently in these cases or did we miss proteins with a low homology in our HMM searches? Interestingly, in kinetoplastids where only Aurora and INCENP (TbCPC1) can be identified by homology searches, a structurally unrelated protein, TbCPC2, has been identified as a new component of the CPC (30, 51). In addition, in the brown alga *Ectocarpus siliculosus*, we find a fusion of the Borealin N-terminus with an FHA domain similar to those found in BORI orthologs of associated lineages (Fig. 8*A*). These changes in CPC composition suggest that a triple helix, as the basis for the CPC scaffold, is not an absolute requirement and alternative conformations may exist.

Albeit centromere concentration of the CPC in mitosis seems to be a general requirement in eukaryotes, the molecular machinery to achieve this displays some plasticity. In this respect, it will be interesting to evaluate the relevance of the Borealin – Shugoshin – H2AT120ph pathway for recruiting the CPC to the centromeres in plants. Notably, *bub1* mutants as well as *sgo1 sgo2* double mutants in Arabidopsis are viable and do not show any significant growth defects (40, 52). Furthermore, Arabidopsis BORIs without a BORR interaction domain still strongly accumulate at centromeres, indicating a prominent role for H3T3^ph^ in recruiting BORIs to centromeres in mitosis while localization of the CPC to the phragmoplast at anaphase is lost. In other organisms, the CPC relocation at anaphase is presumably dependent on the mitotic kinesin MKLP2, a member of the kinesin-6 family (18, 53). Recent study indicates that MKLP2 directly binds to the scaffold proteins of the CPC, and transports the CPC along microtubules to midzone (17). In plants, several kinesins, e.g., NACK1/2, POK1/2, and AtPAKRP1/1L, have been found to accumulate in the midzone of the phragmoplast (54, 55), possibly contributing to CPC relocation.

The acquisition of the phosphate binding BIR domain in animal Survivin likely linked this CPC scaffold protein to an additional, unrelated function. BIR domains were first identified through sequence similarity among the Inhibitors of Apoptosis (IAP) proteins that counteract spontaneous apoptotic programmed cell death (PCD) by interfering with caspase activities. Indeed, most of the BIR-containing proteins, including Survivin, play an inhibitory role in the caspase-dependent apoptotic pathway. Consistent with an absence of apoptosis in plants (28), early members of the plant lineage have acquired a different phospho-binding domain, the FHA domain, to achieve H3T3^ph^ binding. Curiously, an additional unrelated function has also been described for one of the Arabidopsis BORIs, i.e., BORI1, but in this case as a transcriptional repressor of *PEX11b* regulating peroxisome fission (32). DNA binding was suggested to occur via two copies of an AT-hook module, a DNA binding domain that targets AT-rich DNA sequences. Notably, this AT-hook module is only present in the central domain of BORI1 but in BORI2 indicating that it is not required for CPC function.

Taken together, our characterization of BORIs in Arabidopsis led to a redefinition of the minimal architectural requirement for Survivin-type proteins in the context of the CPC, allowing for the identification of putative orthologs in most eukaryotic clades. The here-described case of molecular convergence by differential domain recruitment indicates that we need to broaden our search algorithms in the hunt for orthologs to incorporate such possibilities, and to be more mindful of short helices/motifs as the basis for defining a gene family. Our analyses furthermore illustrate a new example for the contribution of deltaproteobacterial genes to the origin of eukaryotic pathways, lateral gene transfers that have gained increasing interest in recent work (56, 57). Finally, our findings indicate flexibility in the molecular solutions for concentrating the CPC at centromeres in (pro)metaphase found amongst different eukaryotic lineages. These patterns of flexibility and recurrent changes are reminiscent of the rapid molecular evolution often observed for kinetochore and centromere-proximal proteins (20), which are well explained by the paradoxical evolutionary arms race between centromeric DNA and its direct interacting proteins caused by asymmetric meiosis in most eukaryotes, also known as the centromere drive hypothesis (58). Recent work on the mechanisms of centromere drive in mice indeed points towards a delicate balance between regulation of the kinetochore superstructure and the (peri)centromeric chromatin environment, to which Aurora kinase activity has broad regulatory input (59, 60). The patterns of recurrent evolution of the Survivin/*BORI* gene family facilitating the differential recruitment of the CPC to chromatin that we find in our study might thus hallmark evolutionary events that fuel the ongoing war between overzealous centromeres and kinetochores in ancestral eukaryotic lineages.

## Materials and Methods

### Plant Materials and Growth Conditions

The Arabidopsis (*Arabidopsis thaliana*) accession Columbia (Col-0) was used as the wildtype in this study. All mutants are in the Col-0 background. Plants were grown on a solid medium containing half strength Murashige and Skoog salts (MS), 1% (w/v) sucrose, and 1.5% (w/v) agar in a growth chamber at 22°C (16 h of light/8 h of dark). The T-DNA insertion line SALK_095831 (*bori1-1*) was obtained from the Nottingham Arabidopsis Stock Center. The *bori2-1* line was generated by CRISPR/CAS9. Primer pairs for genotyping are described in *SI Appendix*, Table S1 and *SI Appendix*, Fig. S1*A-C*.

### Tandem Affinity Purification (TAP)

Cloning of the transgene encoding a C-terminal GSrhino tag fusion under control of the constitutive cauliflower tobacco mosaic virus 35S promoter and transformation of Arabidopsis cell suspension cultures (PSB-D) was carried out as previously described (31, 61). TAP experiments were performed with 100 mg of total protein extract as input as described in Van Leene et al., 2015. Bound proteins were digested on-bead after a final wash with 500 μL 50 mM NH4HCO3 (pH 8.0). Beads were incubated with 1 μg Trypsin/Lys-C in 50 μL 50 mM NH4OH and incubated at 37°C for 4 h in a thermomixer at 800 rpm. Next, the digest was separated from the beads, an extra 0.5 μg Trypsin/Lys-C was added and the digest was further incubated overnight at 37°C. Finally, the digest was centrifuged at 20800 rcf in an Eppendorf centrifuge for 5 min, the supernatant was transferred to a new 1.5 mL Eppendorf tube, and the peptides were dried in a Speedvac and stored at −20°C until LC-MS/MS analysis. For details on LC-MS/MS and data analysis see *SI Appendix*, Dataset S1.

### Plasmid construction

To create *BORI:GFP* constructs, the genomic fragment of *BORI1* or *BORI2* gene was amplified by PCR and cloned into *pDONR221*. A *Sma*I site was inserted in front of the stop codon of each construct. Both constructs were linearized by *Sma*I digestion and ligated to the monomeric *GFP* (*mGFP*) gene, followed by LR recombination reactions with the destination vector *pGWB501*. To create *BORI_N:GFP* constructs, the C-terminal region of each gene was removed from *BORI:GFP* constructs by inverse PCR. To create *BORI^R3A^:GFP* constructs, corresponding point mutations were introduced in the *BORI:GFP* constructs by inverse PCR. To create the CRISPR/CAS9 construct against *BORI2* gene, the *BORI2* gene-specific spacer sequence was cloned into the *pEn-Chimera*, followed by LR recombination reaction with the destination vector *pDe-CAS9*. To create the amiRNA construct against the *BORI2* gene, 75-bp gene-specific sequences of the BORI2 gene were synthesized and cloned into *pENTR-AtMIR390a-B/c*, followed by LR recombination reactions with the destination vector *pGWB602*. All constructs were transformed into Arabidopsis plants using the floral dip method. Primer pairs for plasmid construction are described in *SI Appendix*, Table S1.

### Production of recombinant BORI proteins

The attR1-attR2 Gateway cassette was amplified by PCR and cloned into *pGEX6p-1*, designated *pGEX6p-GW. BORI1* and *2* cDNAs were amplified from 10-day-old seedling RNA and cloned into *pDONR221* followed by LR recombination reactions with the destination vector *pGEX6p-GW*, and expressed in the *Escherichia coli* strain BL21 (DE3) by induction with 0.1 mM IPTG at 16°C for 16 h. Recombinant proteins were purified by affinity chromatography using Glutathione Sepharose 4B (cytiva) and stored at −80°C.

### Peptide-binding assay

H3^1-21^ and H3^1-21T3ph^ peptides were purchased from AnaSpec, Inc. (Fremont, USA). H3^1-21T11ph^ and H3^1-21T3T11ph^ peptides were synthesized using SCRUM Inc. (Tokyo, Japan). All peptides were biotinylated at the C-terminus. For the peptide-binding assays, 40 pmol protein was incubated with 500 pmol peptide in 200 μl binding buffer (50 mM Tris-HCl, pH 7.5, 150 mM NaCl, 0.1% NP-40) with 0.25% (w/v) BSA at 4°C for 3 h. Streptavidin coated magnetic beads (Dynabeads M-280, Thermo Fisher Scientific) were pre-equilibrated in binding buffer with 0.25% (w/v) BSA. 30 μl beads were added per assay and incubated at 4°C for 1 h. Beads were washed three times with binding buffer and proteins were eluted by boiling in SDS loading buffer. Protein samples were analyzed by SDS-PAGE followed by Western Blotting using 1:2000 diluted anti-GST (Proteintech Group, Inc.) and 1:10000 diluted anti-rabbit IgG, HRP-linked secondary antibody (cytiva).

### Yeast two-hybrid assay

Yeast two-hybrid assays were performed as described in (40). Full length *BORI1* cDNA and the C-terminus of *BORI2* cDNA were amplified by PCR using gene-specific primers from cDNA made from total RNA of wild-type Arabidopsis plants, followed by PCR with universal *attB* primers and cloned into *pDONR221*. The truncated *BORI1* constructs were created by inverse PCR. The subcloned cDNA fragments were recombined into the destination vector *pGBT9* (*DNA-BD*) by LR reaction. Primer pairs for plasmid construction are described in *SI Appendix*, Table S1.

### Embryo observation

Ovules at 4 days after pollination were dissected from siliques and cleared with Herr’s solution: Lactic acid: chloral hydrate: phenol: clove oil: xylene (2:2:2:2:1, w/w), and observed by OLYMPUS BX52 microscopy with differential interference contrast optics.

### Confocal microscopy

For live cell imaging, root tips of 5-day-old seedlings were used. Sample preparation and imaging were performed as described (40).

### Protein extraction and coimmunoprecipitation assay from Arabidopsis seedlings

Coimmunoprecipitation assays were performed as previously described (21). Protein samples were detected with 1:1000 diluted anti-RFP (AB233; Evrogen) for BORR:RFP detection and 1:2000 diluted anti-Histone H3 (ab1791; Abcam) for Histone H3 as primary antibodies and subsequently with 1:10000 diluted anti-rabbit IgG, HRP-linked secondary antibody (cytiva).

### RT-qPCR analysis

RT-qPCR assays were performed as previously described (40). *PP2A3* (AT1G13320) was used as the reference gene. Primer pairs for qPCR are described in *SI Appendix*, Table S1. All experiments were performed in three biological replicates.

### Detection and definition of the BORI/Survivin gene family in eukaryotes

To detect homologs of BORIs and Survivin in eukaryotes, we optimized multiple profile Hidden Markov Models (HMMs) based on iterative reciprocal similarity searches using various tools from the HMMER package version 3.1b2 (62), similar to a strategy used in our previous work (63). Iterative searches with ‘jackhmmer’ were executed with standard inclusion thresholds (E>0.01, bitscore>25) until no new candidate homologs could be included, or as otherwise stated. HMMs were constructed using ‘hmmbuild’ based on multiple sequence alignments of curated homologs. We used both full-length and subdomain (BIR, FHA and helix) HMMs as seeds for reciprocal iterative sequence searches using ‘hmmrsearch’. Our search protocol was based on the following steps/considerations: (I) To limit the amount of homologs to be queried, when searching the widely present BIR and FHA domains and full-length sequences, we used stringent bitscore inclusion cut-offs of 60 up to 70 (--incT 60-70 and --incdomT 60-70). (II) We only considered sequences as candidate Survivin/BORI orthologs if they harbored both a phosphate-binding domain (FHA/BIR) and/or a conserved helix, on the condition that the helix alone should yield reciprocal best hits (phmmer) and/or reciprocal iterative (jackhmmer) hits with bona fide homologs. (III) Putative candidates that contained a single short helical domain were only included in case reciprocal similarity searches yielded phosphate-binding domain containing candidates found in other eukaryotic lineages, and when a particular species or lineage did not yet contain a Survivin/BORI homolog (i.e putative candidates in plants, animals and fungi were excluded). (IV) Were possible, we aimed to optimize one single HMM of the conserved helical domain to capture all Survivin/BORI orthologs (see *SI Appendix*, Dataset S5). We therefore trained an HMM of the gene family-defining feature, the conserved helical domain, on large eukaryotic sequence databases, including our in-house dataset (63), EukProt (64), and UniProt (65). Sequences of orthologs and presence-absence patterns of the Survivin/BORI gene family in a subset of representative eukaryotes can be found in separate text files in *SI Appendix*, Dataset S5. Clade-specific HMMs can be found in *SI Appendix*, Dataset S6.

### Phylogenetic analyses of FHA domains found in BORI-like homologs

To prevent the inclusion of a high number of potential FHA domain-based BORI homologs to consider for phylogenetic analysis, we used a high bitscore cut-off (bitscore>70; see above). Candidate homologs were found in Viridiplantae (Chlorophyta and Streptophyta) and all contained a C-terminal helix (Fig. 1), which strongly suggested that these were orthologous to the BORIs found in Arabidopsis. To find the closest FHA domain to that of BORI, we aligned all Viridiplantae BORI orthologs and generated an HMM of the FHA domain using hmmbuild v3.1b2 (62). Using a similar HMM-based approach, but now with a bitscore cut-off of 60 (--incT 60 --incdomT 60), we found the FHA domains of the PP2C phosphatase KAPP orthologs to be the closest to those of BORI. Subsequent iterations with lower bitscore cut-offs (>50) revealed many non-BORI/KAPP FHA domains to have a roughly similar bitscore, therefore no clear outgroup for BORI and KAPP could be defined in eukaryotes. We therefore searched the Uniprot database for putative prokaryotic homologs. Indeed, Deltaproteobacterial FHA domain-containing sequences were found to be more similar compared to other eukaryotic sequences. To provide an outgroup, we added the seed sequences for the FHA PFAM model (PF00498, see *SI appendix*, Dataset S3). FHA domains were aligned using MAFFT (option g-ins-i) (66). For the phylogenetic analysis in Figure 8C, FHA domain-containing proteins were added that were significantly similar to BORI-like homologs found amongst Stramenopila, Haptista and Cryptista in the EukProt database (64), with a bitscore cut-off (>60). Maximum-likelihood phylogenetic analyses (for analysis files and further description see SI Appendix, Supplementary text, Dataset S3 and Dataset S4) were performed with the IQ-Tree webserver (version 1.6.12) using standard settings for model selection, including the assessment of all mixture models, 1000 Ultrafast bootstrap and SH-like approximate likelihood ratio test replicates (67). Parameters for the final phylogenetic analyses are as follows: analysis Fig. 1*C* (see *SI Appendix*, Dataset S3 – settings: -m LG4X+F -bb 1000 -alrt 1000 -pers 0.3 -numstop 410; 65 positions, 194 sequences with at least 70% occupancy per column, model: LG4X); Fig. 8*C* (see *SI Appendix*, Dataset S4 – settings: -m LG4X+F -bb 1000 -alrt 1000 -pers 0.4 -numstop 300; 64 positions, 408 sequences with at least 70% occupancy per column, model LG4X). Trees were visualized and annotated using FigTree v1.4.4 (68), and/or itol (69).

### Accession Numbers

Sequence data from this article can be found in the Arabidopsis Information Resource database under the following accession numbers: AT2G45490 (AUR3), AT3G02400 (BORI1), AT4G14490 (BORI2), AT4G39630 (BORR), AT5G55820 (INCENP/WYRD), and AT5G19280 (KAPP). Sequences of homologs, and presence-absence patterns of Aurora kinase, INCENP, and Borealin in a subset of representative eukaryotes (see also (20)), can be found in separate text files in *SI Appendix*, Dataset S5.

### Graphics and other software

Plots and alignments were manually compiled into figures using the open-source scalable vector graphics editor Inkscape 1.0rc1 for macOS (Inkscape Project 2020, retrieved from https://inkscape.org). 3D protein structures were visualized using Pymol v2.5. Alignments were manipulated using Jalview (70)

## Supporting information

Komaki_et_al_Supplement

## Acknowledgments

We thank Mariana Motta (University of Hamburg) for critical reading and helpful comments on the manuscript. E. T. is supported by a personal fellowship from the Nederlandse Organisatie voor Wetenschappelijk Onderzoek (grant no. VI.Veni.202.223). This work was supported by the Deutsche Forschungsgemeinschaft (grant no. SCHN 736/8–1) to A.S., and the Japan Society for the Promotion of Science (KAKENHI grant no. JP21K06215) to S.K..

## References

1. M. R. Motta, A. Schnittger, A microtubule perspective on plant cell division. Curr. Biol. 31, R547–R552 (2021).

2. E. C. Tromer, J. J. E. van Hooff, G. J. P. L. Kops, B. Snel, Mosaic origin of the eukaryotic kinetochore. Proc. Natl. Acad. Sci. U. S. A. 116, 12873–12882 (2019).

3. A. Musacchio, A. Desai, A Molecular View of Kinetochore Assembly and Function. Biology 6(2017).

4. S. Müller, G. Jürgens, Plant cytokinesis—No ring, no constriction but centrifugal construction of the partitioning membrane. Semin. Cell Dev. Biol. 53, 10–18 (2016).

5. J. R. Brown, K. K. Koretke, M. L. Birkeland, P. Sanseau, D. R. Patrick, Evolutionary relationships of Aurora kinases: implications for model organism studies and the development of anti-cancer drugs. BMC Evol. Biol. 4, 39 (2004).

6. M. Carmena, M. Wheelock, H. Funabiki, W. C. Earnshaw, The chromosomal passenger complex (CPC): from easy rider to the godfather of mitosis. Nat. Rev. Mol. Cell Biol. 13, 789–803 (2012).

7. A. van der Horst, S. M. A. Lens, Cell division: control of the chromosomal passenger complex in time and space. Chromosoma 123, 25–42 (2014).

8. M. A. Abad, et al., Borealin-nucleosome interaction secures chromosome association of the chromosomal passenger complex. J. Cell Biol. 218, 3912–3925 (2019).

9. F. Wang, et al., A positive feedback loop involving Haspin and Aurora B promotes CPC accumulation at centromeres in mitosis. Curr. Biol. 21, 1061–1069 (2011).

10. F. Wang, et al., Histone H3 Thr-3 phosphorylation by Haspin positions Aurora B at centromeres in mitosis. Science 330, 231–235 (2010).

11. A. E. Kelly, et al., Survivin reads phosphorylated histone H3 threonine 3 to activate the mitotic kinase Aurora B. Science 330, 235–239 (2010).

12. Y. Yamagishi, T. Honda, Y. Tanno, Y. Watanabe, Two histone marks establish the inner centromere and chromosome bi-orientation. Science 330, 239–243 (2010).

13. M. A. Abad, et al., Molecular Basis for CPC-Sgo1 Interaction: Implications for Centromere Localisation and Function of the CPC. bioRxiv, 2021.08.27.457910 (2021).

14. M. A. Hadders, et al., Untangling the contribution of Haspin and Bub1 to Aurora B function during mitosis. J. Cell Biol. 219 (2020).

15. A. J. Broad, K. F. DeLuca, J. G. DeLuca, Aurora B kinase is recruited to multiple discrete kinetochore and centromere regions in human cells. J. Cell Biol. 219 (2020).

16. C. Liang, et al., Centromere-localized Aurora B kinase is required for the fidelity of chromosome segregation. J. Cell Biol. 219 (2020).

17. I. E. Adriaans, et al., MKLP2 Is a Motile Kinesin that Transports the Chromosomal Passenger Complex during Anaphase. Curr. Biol. 30, 2628–2637.e9 (2020).

18. S. Hümmer, T. U. Mayer, Cdk1 negatively regulates midzone localization of the mitotic kinesin Mklp2 and the chromosomal passenger complex. Curr. Biol. 19, 607–612 (2009).

19. O. Kirioukhova, et al., Female gametophytic cell specification and seed development require the function of the putative Arabidopsis INCENP ortholog WYRD. Development 138, 3409–3420 (2011).

20. J. J. van Hooff, E. Tromer, L. M. van Wijk, B. Snel, G. J. Kops, Evolutionary dynamics of the kinetochore network in eukaryotes as revealed by comparative genomics. EMBO Rep. 18, 1559–1571 (2017).

21. S. Komaki, et al., Functional Analysis of the Plant Chromosomal Passenger Complex. Plant Physiol. 183, 1586–1599 (2020).

22. D. Kurihara, S. Matsunaga, T. Omura, T. Higashiyama, K. Fukui, Identification and characterization of plant Haspin kinase as a histone H3 threonine kinase. BMC Plant Biol. 11, 73 (2011).

23. E. Kozgunova, T. Suzuki, M. Ito, T. Higashiyama, D. Kurihara, Haspin has Multiple Functions in the Plant Cell Division Regulatory Network. Plant Cell Physiol. 57, 848–861 (2016).

24. Y. Liu, C. Wang, H. Su, J. A. Birchler, F. Han, Phosphorylation of histone H3 by Haspin regulates chromosome alignment and segregation during mitosis in maize. J. Exp. Bot. 72, 1046–1058 (2021).

25. Q. L. Deveraux, J. C. Reed, IAP family proteins--suppressors of apoptosis. Genes Dev. 13, 239–252 (1999).

26. S. P. Wheatley, D. C. Altieri, Survivin at a glance. J. Cell Sci. 132 (2019).

27. K. Higashi, et al., Identification of a novel gene family, paralogs of inhibitor of apoptosis proteins present in plants, fungi, and animals. Apoptosis 10, 471–480 (2005).

28. A. Daneva, Z. Gao, M. Van Durme, M. K. Nowack, Functions and Regulation of Programmed Cell Death in Plant Development. Annu. Rev. Cell Dev. Biol. 32, 441–468 (2016).

29. E. A. Minina, et al., Apoptosis is not conserved in plants as revealed by critical examination of a model for plant apoptosis-like cell death. BMC Biol. 19, 100 (2021).

30. Z. Li, et al., Identification of a novel chromosomal passenger complex and its unique localization during cytokinesis in Trypanosoma brucei. PLoS One 3, e2354 (2008).

31. J. Van Leene, et al., An improved toolbox to unravel the plant cellular machinery by tandem affinity purification of Arabidopsis protein complexes. Nat. Protoc. 10, 169–187 (2015).

32. M. Desai, R. Pan, J. Hu, Arabidopsis Forkhead-Associated Domain Protein 3 negatively regulates peroxisome division. J. Integr. Plant Biol. 59, 454–458 (2017).

33. D. Durocher, S. P. Jackson, The FHA domain. FEBS Lett. 513, 58–66 (2002).

34. N. Coquelle, J. N. M. Glover, FHA domain pThr binding specificity: it’s all about me. Structure 18, 1549–1550 (2010).

35. A. W. Almawi, L. A. Matthews, A. Guarné, FHA domains: Phosphopeptide binding and beyond. Prog. Biophys. Mol. Biol. 127, 105–110 (2017).

36. J. R. Allen, E. G. Wilkinson, L. C. Strader, Creativity comes from interactions: modules of protein interactions in plants. FEBS J. (2021) https:/doi.org/10.1111/febs.15847.

37. J. M. Stone, M. A. Collinge, R. D. Smith, M. A. Horn, J. C. Walker, Interaction of a protein phosphatase with an Arabidopsis serine-threonine receptor kinase. Science 266, 793–795 (1994).

38. R. W. Williams, J. M. Wilson, E. M. Meyerowitz, A possible role for kinase-associated protein phosphatase in the Arabidopsis CLAVATA1 signaling pathway. Proc. Natl. Acad. Sci. U. S. A. 94, 10467–10472 (1997).

39. I. M. Rienties, J. Vink, J. W. Borst, E. Russinova, S. C. de Vries, The Arabidopsis SERK1 protein interacts with the AAA-ATPase AtCDC48, the 14-3-3 protein GF14lambda and the PP2C phosphatase KAPP. Planta 221, 394–405 (2005).

40. S. Komaki, A. Schnittger, The Spindle Assembly Checkpoint in Arabidopsis Is Rapidly Shut Off during Severe Stress. Dev. Cell 43, 172–185.e5 (2017).

41. I. J. Byeon, S. Yongkiettrakul, M. D. Tsai, Solution structure of the yeast Rad53 FHA2 complexed with a phosphothreonine peptide pTXXL: comparison with the structures of FHA2-pYXL and FHA1-pTXXD complexes. J. Mol. Biol. 314, 577–588 (2001).

42. L. J. Alderwick, et al., Molecular structure of EmbR, a response element of Ser/Thr kinase signaling in Mycobacterium tuberculosis. Proc. Natl. Acad. Sci. U. S. A. 103, 2558–2563 (2006).

43. J. Liu, et al., Structural mechanism of the phosphorylation-dependent dimerization of the MDC1 forkhead-associated domain. Nucleic Acids Res. 40, 3898–3912 (2012).

44. G.-I. Lee, Z. Ding, J. C. Walker, S. R. Van Doren, NMR structure of the forkhead-associated domain from the Arabidopsis receptor kinase-associated protein phosphatase. Proc. Natl. Acad. Sci. U. S. A. 100, 11261–11266 (2003).

45. J. Eswaran, et al., Structure and functional characterization of the atypical human kinase haspin. Proc. Natl. Acad. Sci. U. S. A. 106, 20198–20203 (2009).

46. Y. Lu, et al., Involvement of MEM1 in DNA demethylation in Arabidopsis. Plant Mol. Biol. 102, 307–322 (2020).

47. M. Yamada, G. Goshima, Mitotic Spindle Assembly in Land Plants: Molecules and Mechanisms. Biology 6 (2017).

48. Q. Chen, H. Li, A. De Lozanne, Contractile ring-independent localization of DdINCENP, a protein important for spindle stability and cytokinesis. Mol. Biol. Cell 17, 779–788 (2006).

49. H. Li, et al., Dictyostelium Aurora kinase has properties of both Aurora A and Aurora B kinases. Eukaryot. Cell 7, 894–905 (2008).

50. J. M. Goldberg, et al., The dictyostelium kinome--analysis of the protein kinases from a simple model organism. PLoS Genet. 2, e38 (2006).

51. H. Hu, et al., The Aurora B kinase in Trypanosoma brucei undergoes post-translational modifications and is targeted to various subcellular locations through binding to TbCPC1. Mol. Microbiol. 91, 256–274 (2014).

52. L. Cromer, et al., Centromeric cohesion is protected twice at meiosis, by SHUGOSHINs at anaphase I and by PATRONUS at interkinesis. Curr. Biol. 23, 2090–2099 (2013).

53. U. Gruneberg, R. Neef, R. Honda, E. A. Nigg, F. A. Barr, Relocation of Aurora B from centromeres to the central spindle at the metaphase to anaphase transition requires MKlp2. J. Cell Biol. 166, 167–172 (2004).

54. A. Smertenko, et al., Phragmoplast microtubule dynamics - a game of zones. J. Cell Sci. 131 (2018).

55. A. Herrmann, et al., Dual localized kinesin-12 POK2 plays multiple roles during cell division and interacts with MAP65-3. EMBO Rep. 19 (2018).

56. Y. Hoshino, E. A. Gaucher, Evolution of bacterial steroid biosynthesis and its impact on eukaryogenesis. Proc. Natl. Acad. Sci. U. S. A. 118(2021).

57. P. López-García, D. Moreira, The Syntrophy hypothesis for the origin of eukaryotes revisited. Nat Microbiol 5, 655–667 (2020).

58. S. Henikoff, K. Ahmad, H. S. Malik, The centromere paradox: stable inheritance with rapidly evolving DNA. Science 293, 1098–1102 (2001).

59. T. Kumon, et al., Parallel pathways for recruiting effector proteins determine centromere drive and suppression. Cell 184, 4904–4918.e11 (2021).

60. T. Akera, E. Trimm, M. A. Lampson, Molecular Strategies of Meiotic Cheating by Selfish Centromeres. Cell 178, 1132–1144.e10 (2019).

61. J. Van Leene, et al., Isolation of transcription factor complexes from Arabidopsis cell suspension cultures by tandem affinity purification. Methods Mol. Biol. 754, 195–218 (2011).

62. S. R. Eddy, Accelerated Profile HMM Searches. PLoS Comput. Biol. 7, e1002195 (2011).

63. E. C. Tromer, T. A. Wemyss, P. Ludzia, R. F. Waller, B. Akiyoshi, Repurposing of synaptonemal complex proteins for kinetochores in Kinetoplastida. Open Biol. 11, 210049 (2021).

64. D. J. Richter, C. Berney, J. F. H. Strassert, F. Burki, C. de Vargas, EukProt: a database of genome-scale predicted proteins across the diversity of eukaryotic life. 2020.06.30.180687 (2020).

65. UniProt Consortium, UniProt: a worldwide hub of protein knowledge. Nucleic Acids Res. 47, D506–D515 (2019).

66. K. Katoh, D. M. Standley, MAFFT multiple sequence alignment software version 7: improvements in performance and usability. Mol. Biol. Evol. 30, 772–780 (2013).

67. J. Trifinopoulos, L.-T. Nguyen, A. von Haeseler, B. Q. Minh, W-IQ-TREE: a fast online phylogenetic tool for maximum likelihood analysis. Nucleic Acids Res. 44, W232–5 (2016).

68. A. Rambaut, FigTree v1. 4. Molecular evolution, phylogenetics and epidemiology. Edinburgh, UK: Retrieved from http://tree.bio.ed.ac.uk/software/figtree [Google Scholar] (2012).

69. I. Letunic, P. Bork, Interactive Tree Of Life (iTOL) v4: recent updates and new developments. Nucleic Acids Res. 47, W256–W259 (2019).

70. A. M. Waterhouse, J. B. Procter, D. M. A. Martin, M. Clamp, G. J. Barton, Jalview Version 2--a multiple sequence alignment editor and analysis workbench. Bioinformatics 25, 1189–1191 (2009).

71. F. Burki, A. J. Roger, M. W. Brown, A. G. B. Simpson, The New Tree of Eukaryotes. Trends Ecol. Evol. (2019) https:/doi.org/10.1016/j.tree.2019.08.008.

