## Supplementary material for "Molecular convergence by differential domain acquisition is a hallmark of chromosomal passenger complex evolution": Komaki_et_al_Supplement

**This PDF file includes:**

Figures S1 to S3
Table S1
Legends for Movies S1 to S9
Legends for Datasets S1 to S6

**Other supplementary materials for this manuscript include the following:**

Movies S1 to S9
Datasets S1 to S6

### Supplementary Figures

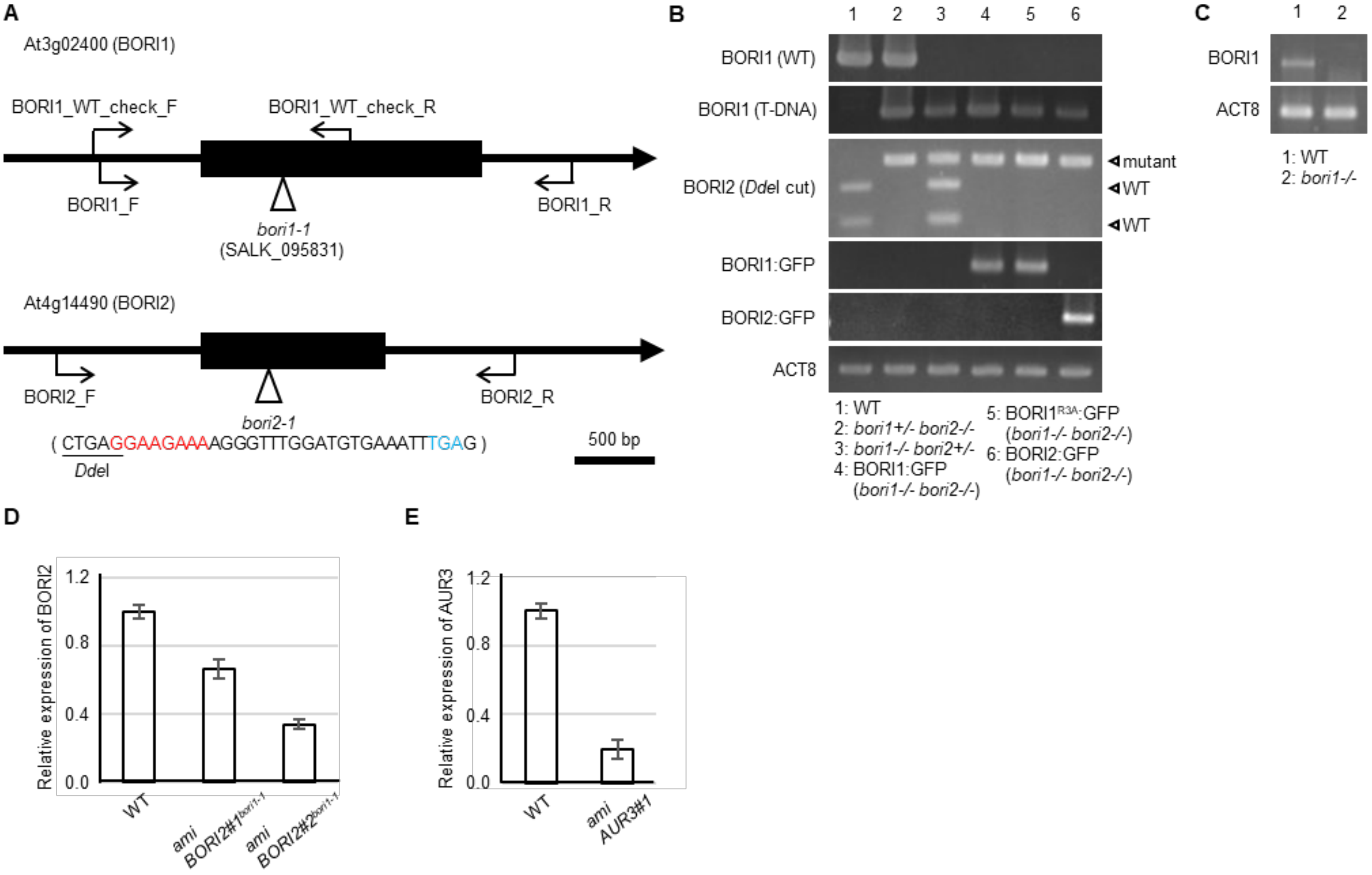


**Figure S1.** Genotyping of *bori* mutants (*A*) Gene structures of *BORI* genes in *Arabidopsis thaliana*. Arrowheads indicate the position of the here-presented mutations. Red color indicates the deleted nucleotides and blue color indicates the premature stop codon in *bori2-1* mutant. Rescue constructs include the region that are amplified by BORI1_F and BORI1_R or BORI2_F and BORI2_R primers. (*B*) Genotyping of *bori* mutants and *BORI* complementation lines by PCR. Only the PCR product from WT BORI2 is digested by the *Dde*I restriction enzyme. ACT8 is used as a control. (*C*) Expression analysis of *BORI1* in *bori1* mutants by RT-PCR using ACT8 as control. (*D*) Relative expression level of *BORI2* in the *amiBORI2* plants was confirmed by RT-qPCR analysis with three biological replicates. Error bars represent means ± SD. (*E*) Relative expression level of *AUR3* in the *amiAUR3* plants was confirmed by RT-qPCR analysis with three biological replicates. Error bars represent means ± SD.

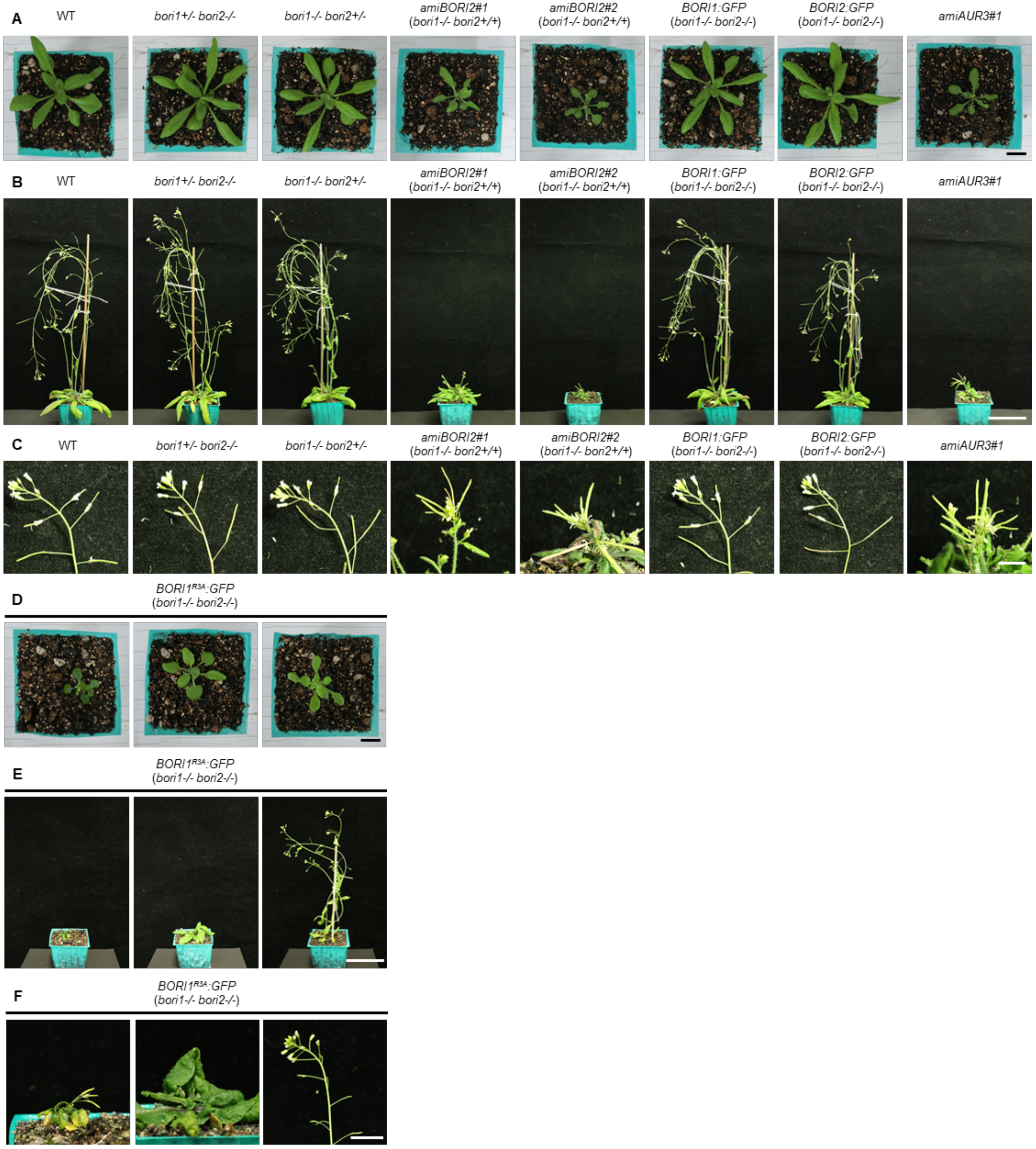
A
WT
bori1+/- bori2-/-
bori1-/- bori2+/-
amiBORI2#1 (bori1-/- bori2+/+)
amiBORI2#2 (bori1-/- bori2+/+)
BORI1:GFP (bori1-/- bori2-/-)
BORI2:GFP (bori1-/- bori2-/-)
amiAUR3#1
B
C
D
BORI1R3A:GFP (bori1-/- bori2-/-)
E
F

**Figure S2.** Phenotypes of *bori* and *aur3* mutants (*A*) 3-week-old plants are shown.Scale bar, 1 cm. (*B*) 30-day-old plants are shown. Scale bar, 5 cm. (*C*) Inflorescences of 5-week-old plants are shown. Scale bar, 1 cm. (*D*) 3-week-old plants expressing $BORI1^{R3A}$*:GFP* are shown Scale bar, 1 cm. (*B*) 30-day-old plants expressing $BORI1^{R3A}$*:GFP* are shown Scale bar, 5 cm. (*C*) Inflorescences of 5-week-old plants expressing $BORI1^{R3A}$*:GFP* are shown Scale bar, 1 cm.

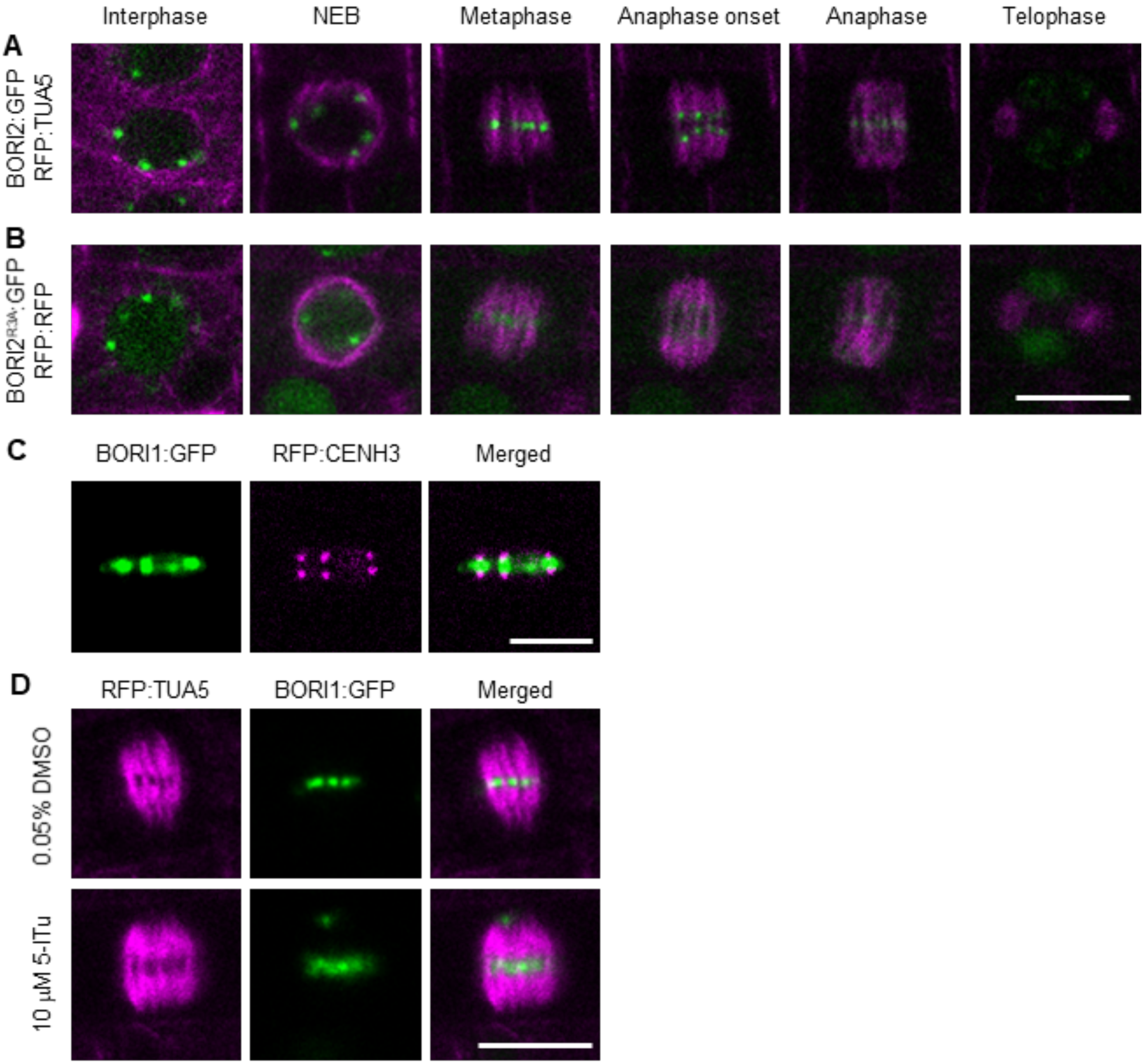


**Figure S3.** Subcellular localization of BORIs. (*A* and *B*) Subcellular localization of BORI2:GFP (*A*) or BORI2$^{R3A}$:GFP (*B*) during the cell cycle. Microtubule structures were visualized by RFP:TUA5. Scale bar, 10 μm. (*C*) Colocalization of BORI1:GFP and inner kinetochore marker RFP:CENH3. Scale bar, 5 μm. (*D*) Subcellular localization of BORI1:GFP in metaphase cells. 5-day-old seedlings were treated without 5-ITu (0.05% DMSO control) or with 10 μM 5-ITu for 90 min. Microtubule structures were visualized by RFP:TUA5. Scale bar, 10 μm.

**Table S1.** Primers used in this study.

| Purpose | Primer name | Sequence (5'-3') | Template |
|---|---|---|---|
| CRISPR | | | |
| CRISPR for BORI2 | BORI2_CRISPR_F | attgGATATCACTGAGGAAGAAAA | |
| | BORI2_CRISPR_R | aaacTTTTCTTCCTCAGTGATATC | |
| Genotyping | | | |
| T-DNA of *bori1-1* | BORI1_T-DNA_check_F | AAAATTCCATTATTTTGAATCAAAGC | |
| | SALK_LBb1.3 | ATTTTGCCGATTTCGGAAC | |
| WT of *bori1-1* | BORI1_WT_check_F | AATGTTGTTTTGCAGGTGGATGAGAAG | |
| | BORI1_WT_check_R | GCTTCATTTTCCTTATCTTCTTCAAC | |
| 1st PCR for *bori2-1* | BORI2_CAPS_F | AATCTCGGAGATGGCGATGTTATC | |
| | BORI2_check_R | TGGACACTTTGTCAATTCAACACATAC | |
| 2nd PCR for *bori2-1* | BORI2_CAPS_F | AATCTCGGAGATGGCGATGTTATC | |
| | BORI2_CAPS_R | CTCTACCTTCTCTACATTAACCAC | |
| BORI1:GFP | BORI1_check_F | GGTTTGATGCCATAGAGGTTCAACTGC | |
| | GFP_R | TTGAAGTCGATGCCCTTCAG | |
| BORI2:GFP | BORI2_check_F | GAGGATCTGGTGGAGAAAAATGGAAGC | |
| | GFP_R | TTGAAGTCGATGCCCTTCAG | |
| ACT8 | ACT8_F | ATGAAGATTAAGGTCGTGGCA | |
| | ACT8_R | TCCGAGTTTGAAGAGGCTAC | |
| RT-PCR | | | |
| BORI1 | BORI1_RT_F | ATGGTTCCGCCATTGCTGAAGCTCGAC | |
| | BORI1_RT_R | TTAGTCGCCCTTTTCCTTTTGCTCTGC | |
| ACT8 | ACT8_RT_F | Same as ACT8_F | |
| | ACT8_RT_R | Same as ACT8_R | |
| qRT-PCR | | | |
| RT-qPCR for BORI2 | BORI2_qRT_F | ATGGTTACGCCATCGTTGAGA | |
| | BORI2_qRT_R | CGAACGATTCGACCAACACG | |
| RT-qPCR for AUR3 | AUR3_qRT_F | GACAGAATCTGACGCTGGCA | |
| | AUR3_qRT_R | CCAAGTGGTCTCCCGATCTC | |
| RT-qPCR for PP2AA3 | PP2AA3_qRT_F | GCCAAAATGATGCAATCTCTC | |
| | PP2AA3_qRT_R | CCGTATCATGTTCTCCACAAC | |
| Cloning of FP fusion constructs | | | |
| Universal *attB* primers | attB1 adapter | GGGGACAAGTTTGTACAAAAAAGCAGGCT | |
| | attB2 adapter | GGGGACCACTTTGTACAAGAAAGCTGGGT | |
| BORI1:GFP | BORI1_F | caaaaaagcaggctccaccGCTGACATGGCAAAGTGGTGAATCTAG | genomic DNA |
| | BORI1_R | caagaaagctgggttGAACAGGAAAACTAAAGCTTCGATACGG | |
| | Cter_SmaI_BORI1_F | gggTAAGTACTAAGTTGATTTGATCCC | pDONR221/BORI1 |
| | Cter_SmaI_BORI1_R | gggGTCGCCCTTTTCCTTTTGCTCTGC | |
| BORI1_N:GFP | BORI1_N:GFP_F | CCCGGGggtggcggtggatcaggcgga | pDONR221/BORI1:GFP |
| | BORI1_N:GFP_R | CTCCAATCCAAAACATTCATTTTC | |
| BORI1$^{R3A}$:GFP | 1st PCR for BORI1$^{R3A}$:GFP_F | gcAGCTGGCGATTCTCTAGGGTTTAAACC | pDONR221/BORI1:GFP |
| | 1st PCR for BORI1$^{R3A}$:GFP_R | TGGACCCTGGGTGAAGTCGAGCTTCAG | |
| | 2nd PCR for BORI1$^{R3A}$:GFP_F | gcTGGTAATGAAATCGCCATCAAAG | 1st PCR product. |
| | 2nd PCR for BORI1$^{R3A}$:GFP_R | GACGATTgcACCAATTCGGATAGTGGATC | |
| BORI2:GFP | BORI2_F | caaaaaagcaggctccaccGATCAAAATAGGTCTCATCCATAGCGG | genomic DNA |
| | BORI2_R | caagaaagctgggttTGAGATTTTACCTTTGGTCTTTCCAGG | |
| | Cter_SmaI_BORI2_F | gggTGAATTGACAAAGTGTCCAACTG | pDONR221/BORI2 |
| | Cter_SmaI_BORI2_R | gggACACATACACGCTTTAGCCTGAAC | |
| BORI2$^{R3A}$:GFP | 1st PCR for BORI2$^{R3A}$:GFP_F | gcAGAAGGCGATGCTCTAGATTACAAG | pDONR221/BORI2:GFP |
| | 1st PCR for BORI2$^{R3A}$:GFP_R | TGGACCCTTGACGAATACTAATCTC | |
| | 2nd PCR for BORI2$^{R3A}$:GFP_F | gcTGGTAATGAAATCGCAATTAAAGACG | 1st PCR product. |
| | 2nd PCR for BORI2$^{R3A}$:GFP_R | AACGATTgcACCAACACGAATCGTAGATC | |
| Histone H3.1:RFP | Histone H3.1_F | GACTGGATCCTGGATCCATCTCTTACTTTCCGTAGATC | genomic DNA |
| | Histone H3.1_R | GTGCGGCCGCAGAAGGTTGAGGTGAAGTCAAAGAGTTG | |
| | pENTR2B for Histone H3.1_F | TCACCTCAACCTTCTGCGGCCGCACTCGAGATATCTAG | pENTR2B |
| | pENTR2B for Histone H3.1_R | TAAGAGATGGATCCAGGATCCAGTCGACTGAATTGGTTC | |
| | Cter_SmaI_ Histone H3.1_F | gggTAAGATTTGAAGAAGATGAAGAAG | pENTR2B/Histone H3.1 |
| | Cter_SmaI_ Histone H3.1_R | gggAGCCCTCTCACCTCTAATCCTCCTC | |
| Yeast two-hybrid assay | | | |
| BORI1_1-585 | BORI1_1-585_F | caaaaaagcaggctccaccATGGTTCCGCCATTGCTGAAGCTCGAC | genomic DNA |

| | | | |
|---|---|---|---|
| | BORI1_1-585_R | caagaaagctgggttTTAGTCGCCCTTTTCCTTTTGCTCTGC | |
| BORI1_1-293 | BORI1_1-293_F | Same as BORI1_1-585_F | genomic DNA |
| | BORI1_1-293_R | caagaaagctgggttTTACTCCAATCCAAAACATTCATTTTC | |
| BORI1_294-585 | BORI1_294-585_F | GTTAAAGATGAGAAGAGAACTACAAGG | pDONR221/BORI1_1-585 |
| | BORI1_294-585_R | CATGGTGGAGCCTGCTTTTTTGTAC | |
| BORI1_535-585 | BORI1_535-585_F | TTAGAAAAAATGAAACTAAGAGAATGG | pDONR221/BORI1_1-585 |
| | BORI1_535-585_R | Same as BORI1_294-585_R | |
| BORI1_294-534 | BORI1_294-534_F | Same as BORI1_1-293_F | pDONR221/BORI1_294-585 |
| | BORI1_294-534_R | TCCATCTTCTCTAACAGTACAGTTCAG | |
| BORI2_287-386 | BORI2_287-386_F | caaaaaagcaggctccaccATGAACAAAGGGAAGAACAAGAAAGCGG | genomic DNA |
| | BORI2_287-386_R | caagaaagctgggttTCAACACATACACGCTTTAGCCTGAAC | |
| Peptide-binding assay | | | |
| BORI1_N | BORI1_N_F | Same as BORI1_1-585_F | genomic DNA |
| | BORI1_N_R | Same as BORI1_1-293_R | |
| BORI1_$N^{R3A}$ | BORI1_$N^{R3A}$_F | Same as BORI1_1-585_F | pDONR221/BORI1$^{R3A}$:GFP |
| | BORI1_$N^{R3A}$_R | Same as BORI1_1-293_R | |
| BORI2_N | BORI2_N_F | caaaaaagcaggctccaccATGGTTACGCCATCGTTGAGATTAGTATTC | genomic DNA |
| | BORI2_N_R | caagaaagctgggttTCACTCTACCTTCTCTACATTAACCAC | |
| BORI2_$N^{R3A}$ | BORI2_$N^{R3A}$_F | Same as BORI2_N_F | pDONR221/BORI2$^{R3A}$:GFP |
| | BORI2_$N^{R3A}$_R | Same as BORI2_N_R | |
| artificial microRNA | | | |
| amiBORI2 | amiBORI2_F | TGTATTGTAATCTAGAGCATCGCCAATGATGATCACATTCGTTATCTATTTTTTTGGCGATGCTATAGATTAC AA | |
| | amiBORI2_F | AATGTTGTAATCTATAGCATCGCCAAAAAAATAGATAACGAATGTGATCATCATTGGCGATGCTCTAGATTAC AA | |
| amiAUR3 | amiAUR3_F | TGTATTCAGCGCCACTATGTACCTCATGATGATCACATTCGTTATCTATTTTTTGAGGTACATATTGGCGCTG AA | |
| | amiAUR3_R | AATGTTCAGCGCCAATATGTACCTCAAAAAATAGATAACGAATGTGATCATCATGAGGTACATAGTGGCGCT GAA | |

### Supplementary Movies

All SI Movies can also be found under the following FigShare link:
https://figshare.com/s/c5d3044493e48b310fb7

**Movie S1**

Subcellular localization of BORI1:GFP during mitosis.

**Movie S2**

Subcellular localization of BORI2:GFP during mitosis.

**Movie S3**

Co-localization of BORI1:GFP and BORR:RFP during mitosis.

**Movie S4**

Subcellular localization of BORI1_N:GFP during mitosis.

**Movie S5**

Subcellular localization of BORI1$^{R3A}$:GFP during mitosis.

**Movie S6**

Subcellular localization of BORI2$^{R3A}$:GFP during mitosis.

**Movie S7**

3D projection of *bori1 bori2* root cells complemented with *BORI1:GFP*. Both cells have 10-dotted-GFP signals. Left cell is shown in Fig. 7*E*.

**Movie S8**

3D projection of *bori1 bori2* root cells complemented with *BORI1$^{R3A}$:GFP*. The cell has 9-dotted-GFP signals.

**Movie S9**

3D projection of *bori1 bori2* root cells complemented with *BORI1$^{R3A}$:GFP*. Both cells have 11-dotted-GFP signals. Left cell is shown in Fig. 7*F*.

**Dataset S1**

Protein Identification details obtained by mass spectrometry.

**Dataset S2**

Separate .pdb files of predicted 3D structures of AthaBORI 1 and 2, and a pymol session file with BORI1 and 2 aligned corresponding to the structures shown in Figure 1D. BORI1: At3g02400 (uniprot-ID:Q9M8A0); BORI2: At4g14490 (uniprot-ID:O23305) - downloaded from the AF2 repository https://alphafold.ebi.ac.uk/.

**Dataset S3**

Text files and multiple sequence alignment used for IQ-Tree maximum-likelihood phylogenetic analysis found in Figure 1D.

**Dataset S4**

Text files and multiple sequence alignment used for IQ-Tree maximum-likelihood phylogenetic analysis found in Figure 8C.

**Dataset S5**

Text files, including: (1) .xlsx table with genome/transcriptome sources, presences and absences of Aurora kinase, INCENP, Borealin and Survivin/BORI, (2) sequences of homologs in separate text files, and (3) including separate domain annotations for the Survivin/BORI gene family + Hidden Markov models used for these annotations.

**Dataset S6**

Hidden Markov Models and multiple sequence alignment files for the detection of both N- and C-terminal helices in Survivin/BORI orthologs (EukProt sequence database).
